# Trisomy 21 induces pericentrosomal crowding disrupting early stages of primary ciliogenesis and mouse cerebellar development

**DOI:** 10.1101/2021.11.10.468107

**Authors:** Cayla E Jewett, Bailey L McCurdy, Eileen T O’Toole, Katherine S Given, Carrie H Lin, Valerie Olsen, Whitney Martin, Laura G Reinholdt, Joaquin M Espinosa, Kelly D Sullivan, Wendy B Macklin, Rytis Prekeris, Chad G Pearson

## Abstract

Primary cilia are signaling organelles essential for development and homeostasis. Loss of primary cilia is lethal, and decreased or defective cilia cause multisystemic conditions called ciliopathies. Down syndrome shares clinical overlap with ciliopathies. We previously showed that trisomy 21 diminishes primary cilia formation and function due to elevated Pericentrin, a centrosome protein encoded on chromosome 21. Pericentrin is mislocalized, creating aggregates that disrupt pericentrosomal trafficking and microtubule organization. Here, we examine the cilia-related molecules and pathways disrupted in trisomy 21 and their *in vivo* phenotypic relevance. Utilizing ciliogenesis time course experiments, we reveal how Pericentrin, microtubule networks, and components of ciliary vesicles are reorganized for ciliogenesis in euploid cells. Early in ciliogenesis, chromosome 21 polyploidy results in elevated Pericentrin and microtubule networks away from the centrosome that ensnare MyosinVA and EHD1, blocking mother centriole uncapping that is essential for ciliogenesis. Ciliated trisomy 21 cells have persistent trafficking defects that reduce transition zone protein localization, which is critical for Sonic hedgehog signaling. Sonic hedgehog signaling is decreased and anticorrelates with Pericentrin levels in trisomy 21 primary mouse embryonic fibroblasts. Finally, we observe decreased ciliation *in vivo*. A mouse model of Down syndrome with elevated Pericentrin has fewer primary cilia in cerebellar granule neuron progenitors and thinner external granular layers. Our work reveals that elevated Pericentrin in trisomy 21 disrupts multiple early steps of ciliogenesis and creates persistent trafficking defects in ciliated cells. This pericentrosomal crowding results in signaling defects consistent with the neurological deficits found in individuals with Down syndrome.

## Introduction

Trisomy 21 or Down syndrome (DS) is a common chromosomal disorder characterized by phenotypes including craniofacial abnormalities, intellectual disability, heart defects, and cerebellar hypoplasia.^1–3^ These pathologies overlap with those of ciliopathies—genetic disorders affecting primary cilia.^4^ Primary cilia are concentrated signaling hubs, particularly for the Sonic hedgehog pathway (Shh), and are essential for vertebrate development. The primary cilium nucleates from the centrosome and projects into the extracellular space. The centrosome is comprised of a mother and daughter centriole and surrounding pericentriolar material. Cilia formation requires the spatial and temporal coordination of many molecules. It begins with remodeling the mother centriole, including addition of appendages that serve as docking platforms for cargo delivery.^5–7^ This triggers removal and proteasomal degradation of the mother centriole capping proteins, allowing extension of axoneme microtubules and membrane remodeling to ensheath the axoneme with a ciliary membrane.^5,8,9^ A barrier complex forms at the base of the cilium called the transition zone, that restricts access to and from the cilium. This creates a unique ciliary compartment that is biochemically distinct from the rest of the cell, allowing the cilium to function as a specialized signaling organelle.^10^

Building and maintaining signaling cilia requires trafficking of molecules to and from the centrosome.^11,12^ Molecules are trafficked in both membrane-derived vesicles and granular moieties, called centriolar satellites, that move along microtubules nucleated and organized by the centrosome.^13–15^ Pericentrin (PCNT) is an essential centrosome scaffolding protein that, together with CDK5RAP2/CEP215, organizes γ-tubulin and microtubules,^16,17^ and in *Drosophila* the PCNT ortholog is required for cilia function in sensory cells and sperm.^18^ *PCNT* is encoded on human chromosome 21 (HSA21), and we previously showed that elevated PCNT due to increased copy number in trisomy 21 is necessary and sufficient to initiate cilia defects in human DS-derived fibroblasts compared to age- and sex-matched controls.^19^ Elevated PCNT forms aggregates that colocalize with satellite proteins such as PCM1 and disrupts the flux of intracellular components to and from the centrosome, analogous to a traffic jam.^20^ While we showed reduced centrosome localization of the ciliary IFT20 protein in trisomy 21 cells,^19^ IFT20 recruitment to the cilium is a later step in the process of ciliogenesis^9,21^. Thus, it is unknown which molecules or pathways are disrupted in trisomy 21 that reduce cilia formation and signaling, and whether these cilia defects observed in cultured cells contribute to DS-associated phenotypes *in vivo*.

Here, we use isogenic human cell lines to eliminate genetic variability and mouse models of DS to show that elevated PCNT induces trafficking defects around the centrosome such that cargo trafficking events required for early steps in primary ciliogenesis, including ciliary vesicle formation and mother centriole uncapping are held up in a ‘pericentrosomal crowd’. Of the trisomy 21 cells that do ciliate, intracellular trafficking defects persist as transition zone proteins are unable to reach the centrosome efficiently, thereby decreasing their localization at the transition zone. Consistent with transition zone defects, Shh signaling is reduced and anticorrelates with PCNT levels. A mouse model of DS with increased *Pcnt* copy number and elevated PCNT levels has reduced primary cilia in both primary mouse embryonic fibroblasts (MEFs) as well as cerebellar neuronal precursor cells *in vivo*. Consistent with ciliary assembly and signaling defects, these mice have a thinner external granular layer and fewer neuronal protrusions. Our findings reveal how early events in ciliogenesis are disrupted by a PCNT-overexpression-induced crowding phenotype and that these ciliation and signaling defects have consequences for *in vivo* brain development in DS.

## Results

### Rapid PCNT and microtubule reorganization in response to ciliation cues increases with HSA21 dosage

We previously showed that elevated PCNT is necessary and sufficient for decreased ciliation in trisomy 21 cells.^19,20^ Whereas PCNT normally nucleates and organizes microtubules emanating from centrosomes, elevated PCNT has three major consequences in trisomy 21 cells (Figure 1A): 1) PCNT nucleates excess microtubules, thereby increasing microtubule density around the centrosome; 2) PCNT forms large protein aggregates along these microtubules; and 3) PCNT nucleates cytoplasmic microtubules that are disconnected from the centrosome.^19,20^ Together, elevated PCNT perturbs trafficking to and from centrosomes by altering microtubule networks and molecular composition of the pericentrosomal region, thereby hindering the exchange of molecules required for ciliogenesis - pericentrosomal crowding. This model was derived from an analysis of either cycling cells or cells 24 hours after serum depletion (0.5% serum in DMEM) to induce G1 cell cycle arrest and ciliogenesis.^19,20^ We questioned whether elevated PCNT results in an immediate trafficking delay upon induction of ciliogenesis or if trafficking defects build up over time such that only later steps of ciliogenesis are affected.

**Figure 1.**
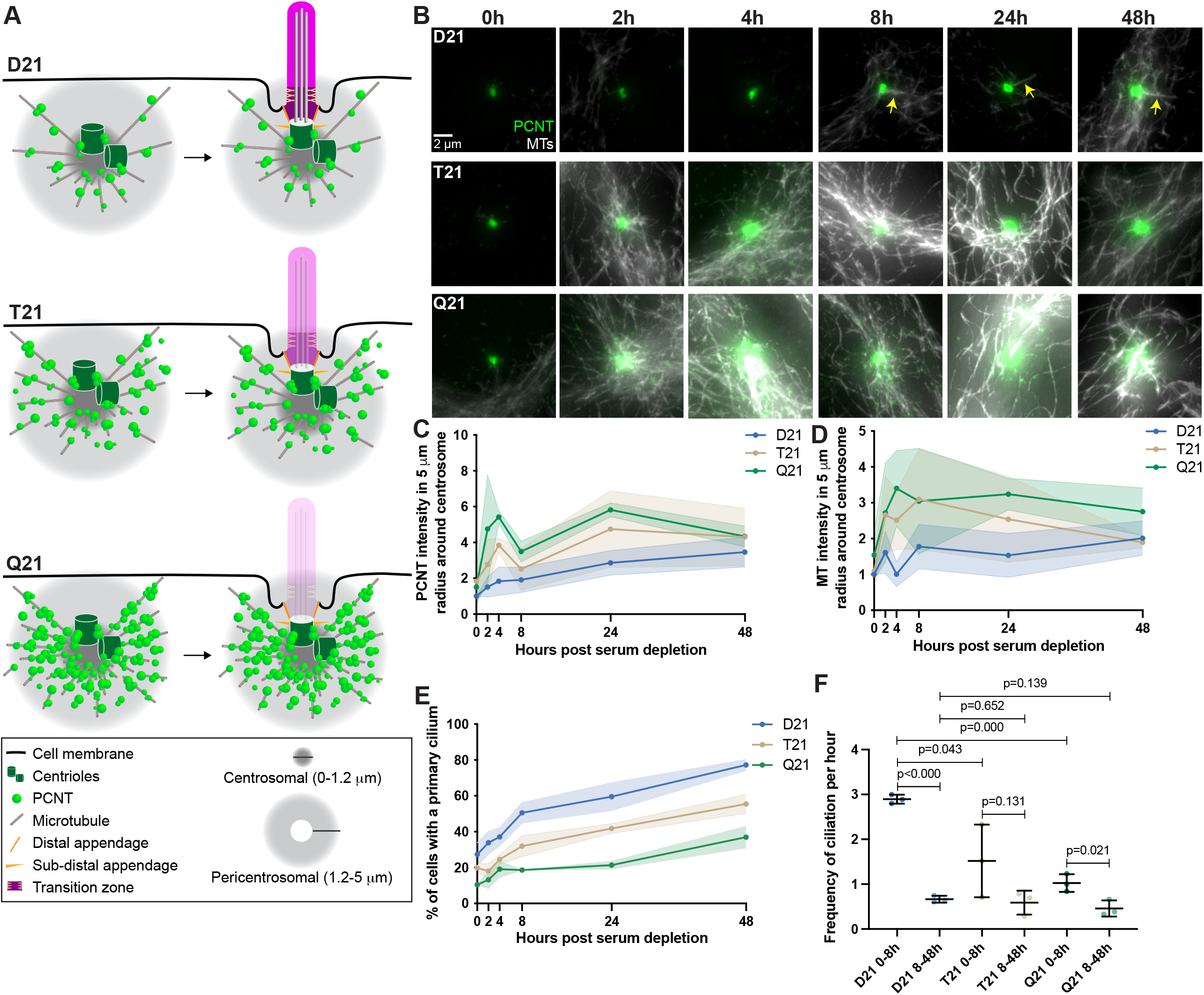
Rapid PCNT and microtubule reorganization in response to ciliation cues increases with HSA21 dosage. (A) Model figure showing the three main consequences of elevated PCNT in trisomy 21 (T21) and tetrasomy 21 (Q21) compared to disomy 21 (D21) cells: 1) PCNT nucleates more microtubules; 2) PCNT forms large protein aggregates on microtubules; and 3) PCNT nucleates microtubules further away from the centrosome. These changes occur predominantly in the pericentrosomal region defined as 1.2-5 μm from the centroid of the centrosome. (B) Representative confocal images from time course experiments of RPE1 D21, T21, and Q21 cells grown on coverslips and serum depleted for 0, 2, 4, 8, 24, and 48 h. Cells were stained with DM1a to label microtubules (MTs) and PCNT. Arrows point to cilium labeled by DM1a staining. (C) Quantitation of PCNT intensities in a 5 μm radial circle around the centrosome throughout the time course normalized to D21 average at 0 h. Graph shows mean ± SD. (D) Quantitation of microtubule intensities in a 5 μm radial circle around the centrosome throughout the time course normalized to D21 average at 0 h. Graph shows mean ± SD. (E) Quantitation of ciliation frequency throughout the time course using DM1a as a marker for cilia. Graph shows mean ± SD. N’s and statistical tests are listed in Table S1.

To differentiate between these alternatives, we used human Retinal Pigmented Epithelial (RPE1) cells to perform a time course during ciliogenesis and analyzed how PCNT and microtubule organization change at the centrosome. In RPE1 cells with two copies of HSA21 (D21), PCNT levels increased rapidly by approximately 50% near the centrosome (within a 5 μm radius) 2 hours after serum depletion, and continued to gradually increase throughout the remaining 48-hour time course (Figures 1B, 1C). Similarly, microtubule density around the centrosome fluctuated between 0 and 8 hours post serum depletion, then remained constant through the rest of the time course (Figures 1B, 1D). Isogenic human RPE1 cells genetically engineered to have three or four copies of HSA21 (Trisomy 21/T21 or Tetrasomy 21/Q21), have elevated PCNT prior to serum depletion both at and around the centrosome (within a 5 μm radius) (Figures 1B, 1C). Microtubule intensity was more similar in all three cell lines prior to serum depletion (Figures 1B, 1D). By 2 hours, both PCNT and microtubule intensities elevated with increasing HSA21 dosage (Figures 1B-D). Elevated PCNT near the centrosome in T21 and Q21 cells persisted through the time course and was not due to changes in whole cell protein levels (Figures 1B-D, S1A-C). Instead, PCNT and microtubules in T21 and Q21 cells reorganized at the centrosome (0.0-1.2 μm region from the centroid of the centrosome) and pericentrosomal region (1.2-5.0 μm region from the centroid of the centrosome), such that more PCNT foci and microtubules were distributed around the centrosome (Figures 1B-D, S1D-G). These data support a model whereby PCNT accumulates at and around the centrosome upon induction of ciliogenesis. Interestingly, by 48 hours, PCNT and microtubule intensities were least changed between the three cell populations (Figures 1B-D, S1D-G), suggesting that T21 and Q21 cells might adapt to an elevated PCNT state upon prolonged G1 arrest.

To understand the interplay between PCNT levels, microtubules, and ciliation, we quantified primary cilia frequency through the time course. D21 cells demonstrated two rates of ciliation: a fast phase from 0-8 hours and a slow phase from 8-48 hours (Figures 1E, 1F). In contrast, T21 and Q21 cells showed a decreased initial fast phase; however, the slow phase from 8-48 hours was similar to D21 cells (Figures 1E, 1F). The fast ciliation phase correlates with the early increases to PCNT and microtubule intensities which are more robust in T21 and Q21 cells (Figures 1B-F). The delay in ciliogenesis observed in T21 and Q21 cells is consistent with a model whereby increasing HSA21 dosage disrupts pericentrosomal trafficking flux to and from the centrosome early in the process of ciliogenesis.

### HSA21 ploidy does not affect centriole appendages but decreases vesicles at the mother centriole

Primary ciliogenesis requires coordination between a series of trafficking and complex assembly events over several hours.^22^ Centriole appendage assembly at the distal end of the mother centriole is an initiating event in ciliogenesis. These appendages serve as a scaffold for receiving ciliary components trafficked to the centrosome.^5-7^ CEP164 and CEP83 are two distal appendage proteins, with CEP83 adjacent to the centriole microtubule walls and CEP164 at the tip of the appendage structure.^23^ In all cell lines (D21, T21, and Q21), CEP164 and CEP83 localized normally at the mother centriole (Figures 2A, S2A). Moreover, the subdistal appendage proteins ODF2^24,25^ and Ninein^26^ correctly localized to the mother centriole (Figures 2B, S2B). Thus, HSA21 ploidy does not affect mother centriole appendages.

**Figure 2.**
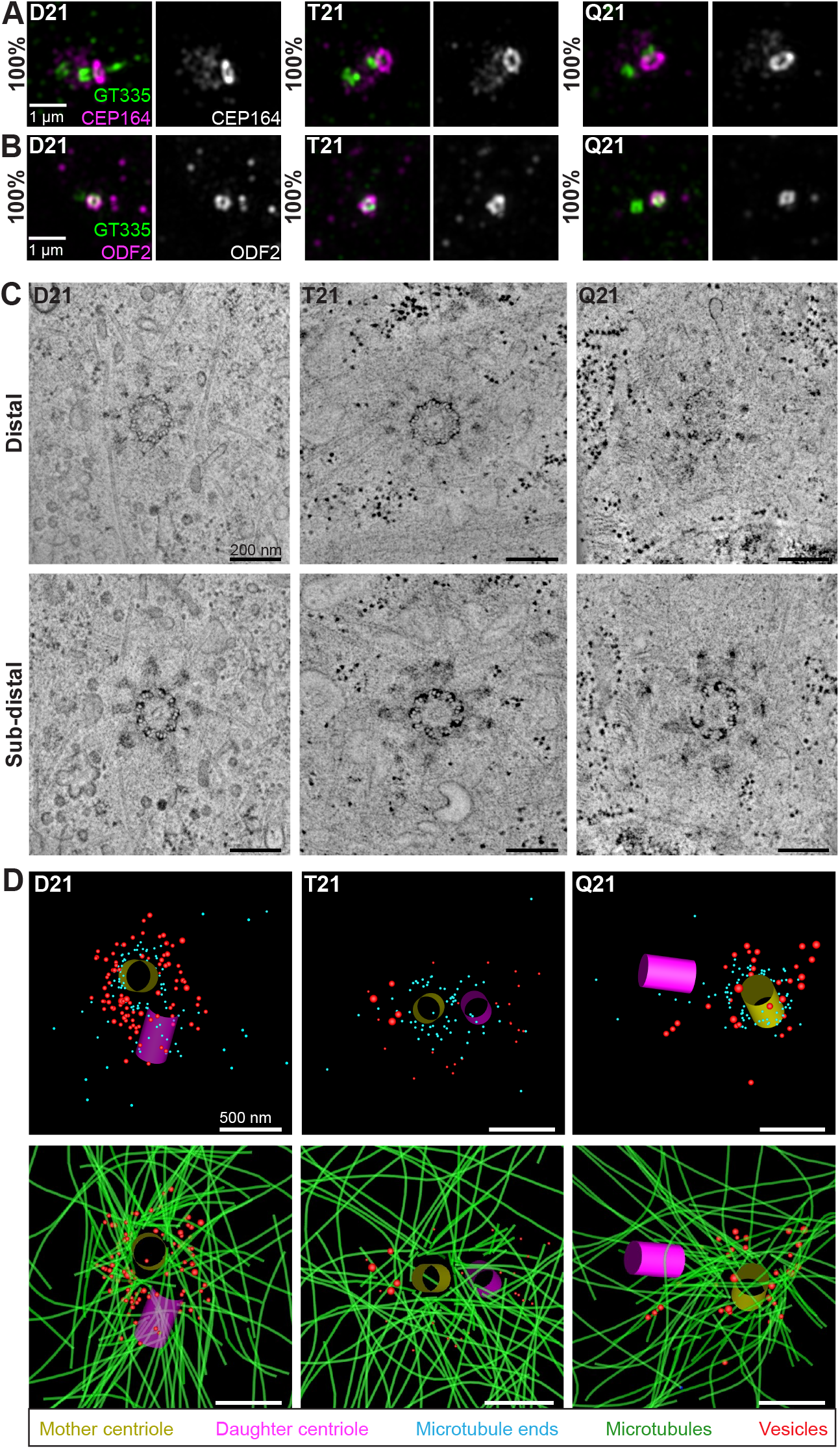
HSA21 ploidy does not affect centriole appendages but decreases vesicles at the mother centriole. (A-B) Representative SIM images of RPE1 D21, T21, and Q21 cells grown on coverslips and serum depleted for 24 h. Cells were stained with GT335 and the distal appendage marker CEP164 (A) or the subdistal appendage marker ODF2 (B). Percentages represent cells with indicated marker for 3 N’s. (C) EM tomograms of RPE1 D21, T21, and Q21cells serum depleted for 24h showing microtubule triplets and distal appendages (top panel) and subdistal appendages (bottom panel). (D) 3D models of EM tomograms at centrosome. Top row shows mother centriole (yellow), daughter centriole (magenta), microtubule minus ends (light blue spheres), and vesicles (red). Bottom row shows models with microtubules (green) and vesicles displayed. Tomograms and models shown are of cells prior to ciliary vesicle formation.

To gain further insight into the ultrastructure of centrioles with increasing HSA21 dosage, we performed 3D electron tomography. Consistent with the immunofluorescence data, D21, T21, and Q21 cells showed ninefold symmetry of distal and subdistal appendages. Moreover, the triplet microtubules that comprise the centriole wall were unchanged across all cell lines, further confirming that HSA21 ploidy does not affect centriole structure. However, 3D modeling of the tomograms revealed changes in the number of vesicles at the mother centriole. D21 cells showed many small vesicles (35-65 nm diameter) near the mother centriole (91 vesicles), whereas vesicle number was decreased in T21 and Q21 cells (26 and 31 vesicles, respectively) (Figure 2D, red spheres). Moreover, in D21 cells, vesicles were distributed along microtubules and at their ends while vesicles in the T21 and Q21 cells were not always found on microtubules (Figure 2D, Video S1-3). Despite the increased microtubule density observed in Figure 1, the comparable number of microtubule minus ends near the centrioles (86, 69, 90 microtubule ends in D21, T21, Q21, respectively) suggests that the additional microtubules are not arising from the mother centriole. Because the tomograms only capture the centrosomal region (0-1.2 μm from centroid of centrosome), this is consistent with our studies showing that more microtubules are found in the pericentrosomal region that is distal to the centrioles (Figure 1D).^20^ Together, these data suggest that HSA21 dosage does not affect centriole structure nor centriole appendage formation but may alter membrane structures at and around the centrosome.

### Preciliary vesicle components contribute to PCNT-induced pericentrosomal crowding

Centriolar appendages are docking sites for delivery of vesicles and molecules required for ciliogenesis.^5^ Our EM tomograms suggested that vesicle accumulation at the mother centriole is defective with HSA21 ploidy. Because increased HSA21 ploidy resulted in changes to the pericentrosomal region in early ciliogenesis, one potential explanation for decreased mother centriole vesicles is disrupted trafficking in the pericentrosomal region. We thus examined whether molecules required for initiating ciliogenesis are disrupted in this region. The first molecules that initiate the downstream steps of ciliogenesis are termed the preciliary or distal appendage vesicle and include the motor protein Myosin VA (MYOVA)^8^ and the membrane shaping protein EHD1.^9^ At 2 hours post serum depletion, about half of D21 cells have formed a MYOVA vesicle at the mother centriole (Figure 3A). In contrast, T21 and Q21 cells show decreased MYOVA vesicle formation and a striking buildup of MYOVA protein in the pericentrosomal region surrounding the centrosome (Figures 3A-B). These changes in pericentrosomal MYOVA intensity were not due to changes in whole cell MYOVA protein levels (Figure S3A). Radial analysis of MYOVA intensity surrounding the centrosome in D21 cells showed high MYOVA levels at the centrosome that then decreased in intensity moving away from the centrosome (Figure 3B inset). We then compared MYOVA intensity distribution at and around the centrosome with increasing ploidy through the time course. Two hours after serum depletion, MYOVA intensities in T21 and Q21 cells were increased in the pericentrosomal 1.2-5.0 μm region from the centroid of the centrosome (Figure 3B). Moreover, centrosomal MYOVA levels were decreased with increasing ploidy (Figure S3B). By 4 hours, pericentrosomal MYOVA intensities in T21 and Q21 cells became more prominent, while MYOVA centrosomal levels remained decreased (Figures 3A, 3C, S3C). By 24 hours, T21 and Q21 cells still showed changes in MYOVA intensity distribution, but the increased intensities were tighter around the centrosome (Figures 3A, 3D, S3D), consistent with the redistribution of PCNT intensities observed between 4 and 24 hours (Figure 1C). Whereas MYOVA intensities increased at the centrosome in T21 and Q21 cells 24 hours post serum depletion, MYOVA appeared more diffuse at the mother centriole compared to D21 cells (Figure 3A, bottom panel). This suggests an additional defect in vesicle coalescence. Taken together, MYOVA intensity increases in the pericentrosomal region with increasing ploidy, and we define this pericentrin-induced redistribution of molecules around the centrosome with increasing HSA21 ploidy as pericentrosomal crowding.

**Figure 3.**
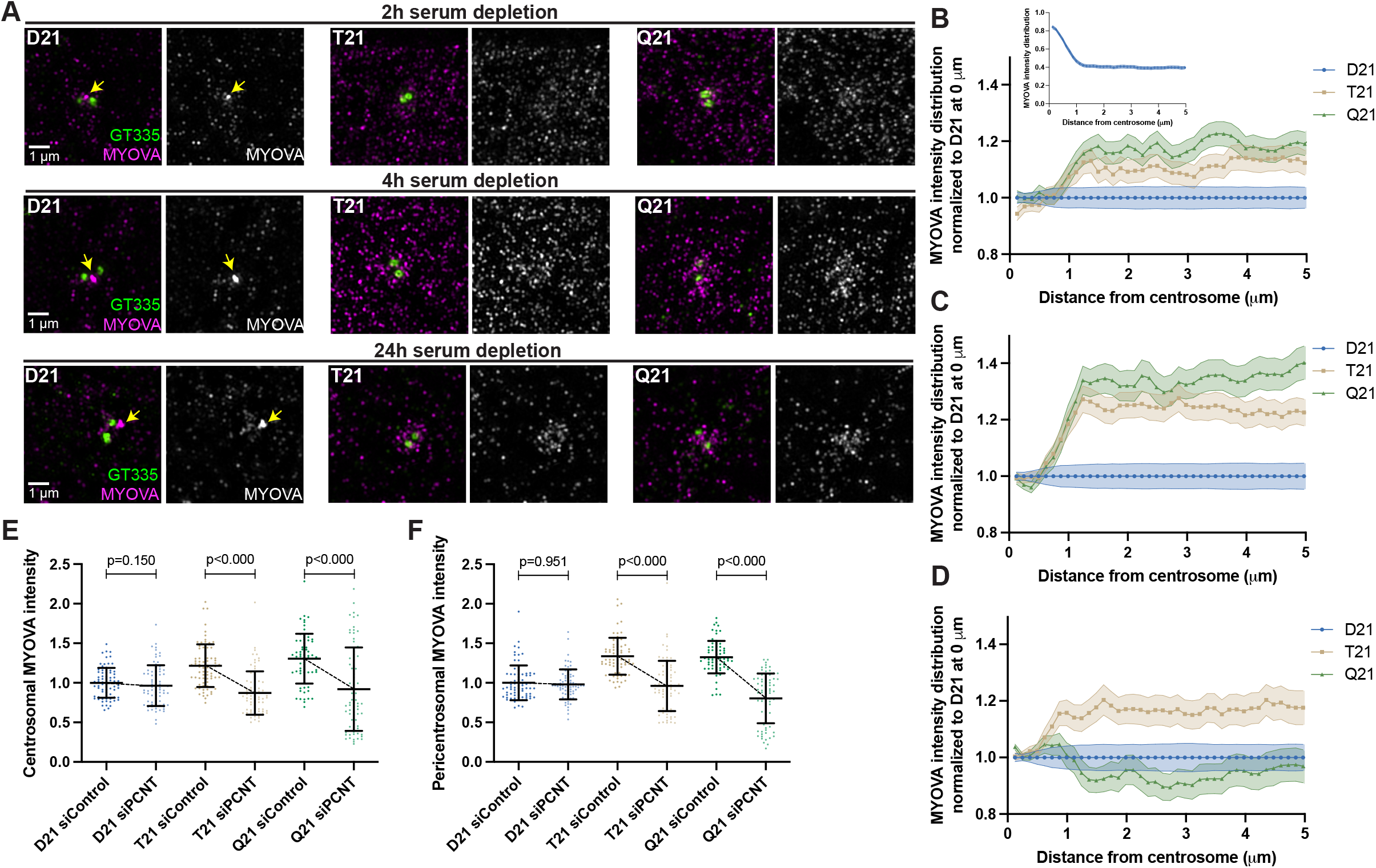
Preciliary vesicle components contribute to PCNT-induced pericentrosomal crowding. (A) Representative structured illumination microscopy (SIM) images from time course experiments of RPE1 D21, T21, and Q21 cells grown on coverslips and serum depleted for 2, 4, and 24 h. Cells were stained with GT335 to label centrioles and MYOVA. (B-D) Distribution of MYOVA intensities moving away from the centrosome for 2 (B), 4 (C), and 24 (D) h timepoints. (E) Quantitation of centrosomal MYOVA intensity in 0-1.2 μm region around centrosome for control and siPCNT treated D21, T21, and Q21 cells. (F) Quantitation of pericentrosomal MYOVA intensity in 1.2-5 μm region around centrosome for control and siPCNT treated D21, T21, and Q21 cells. All values were normalized to the D21 average, and graphs show mean ± SD. N’s and statistical tests are listed in Table S1.

We then asked whether elevated PCNT found with increased HSA21 ploidy was sufficient to increase MYOVA in the pericentrosomal crowding region. We have previously shown that reducing PCNT levels in T21 and Q21 cells with siRNA rescues ciliation.^20^ Reducing PCNT levels in T21 and Q21 cells to D21 levels with siRNA at 24 hours post serum depletion rescued the increased centrosomal and pericentrosomal MYOVA intensity to D21 levels (Figures 3E, 3F, S3E, S3F). Moreover, the pericentrosomal increase in MYOVA consistently increases with increasing PCNT levels (Figures S3G, S3H). Thus, elevated PCNT from increasing HSA21 ploidy induces pericentrosomal crowding where MYOVA accumulates during early ciliogenesis, thereby preventing efficient MYOVA vesicle formation at the mother centriole.

We next examined EHD1, an early vesicle protein that is recruited to the mother centriole for coalescence and fusion of preciliary or distal appendage vesicles.^9^ We observed decreased EHD1 accumulation at the mother centriole with increasing ploidy and a slight increase in EHD1 that is caught up in the pericentrosomal region (Figure S3I). Together, our data support a model whereby elevated PCNT in trisomy 21 accumulates at and around the centrosome immediately after induction of ciliogenesis and induces pericentrosomal crowding, disrupting multiple trafficking pathways required for early preciliary vesicle formation.

### Increased HSA21 ploidy disrupts mother centriole uncapping in a PCNT-dosage dependent manner

Delivery of preciliary vesicles occurs just prior to or coincident with mother centriole uncapping,^8^ which involves removal and degradation of the centriole capping proteins CP110 and CEP97.^27,28^ Both CP110 and CEP97 removal from mother centrioles is defective in T21 and Q21 cells compared to D21 cells (Figures 4A, S4A-C). To determine if these uncapping defects were due to elevated PCNT levels, we decreased PCNT levels in T21 and Q21 cells back to D21 levels and examined uncapping. Strikingly, reduced PCNT levels rescued CP110 removal from the mother centriole (Figure 4B). Together, this indicates that elevated PCNT from increased HSA21 ploidy is sufficient to block mother centriole uncapping. Because uncapping requires the trafficking of the preciliary vesicle, we propose that elevated PCNT-induced crowding ensnares components required for efficient uncapping (Figure 4E).

**Figure 4.**
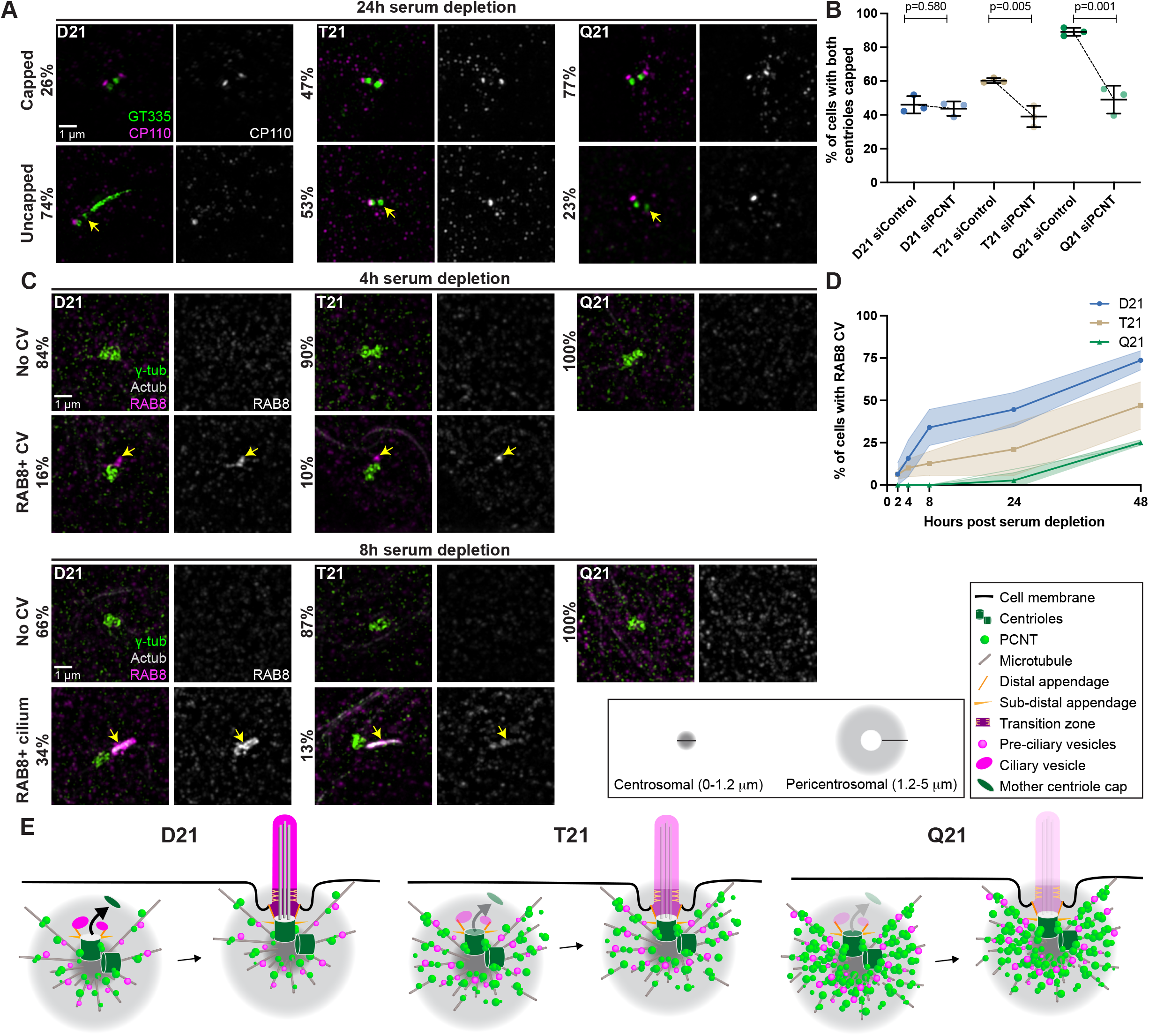
Increased HSA21 ploidy disrupts mother centriole uncapping in a PCNT-dosage dependent manner. (A) Representative SIM images of RPE1 D21, T21, and Q21 cells grown on coverslips and serum depleted for 24 h. Cells were stained with GT335 and the centriole capping protein CP110. Percentages represent cells with indicated phenotype across 3 N’s. (B) Quantitation of CP110 centriole capping for control and siPCNT treated D21, T21, and Q21 cells. (C) Representative SIM images of RPE1 D21, T21, and Q21 cells grown on coverslips and serum depleted for 4, and 8 h. Cells were stained with γ-tubulin, Actub, and RAB8. Percentages represent cells with indicated phenotype across 3 N’s. All time points are shown in Figure S3D. (D) Quantitation of cells with RAB8-positive ciliary vesicle throughout the time course. (E) Cartoon model depicting altered distribution of MYOVA from the mother centriole to the pericentrosomal region and decreased mother centriole uncapping with increasing ploidy. Graphs shows mean ± SD. N’s and statistical tests are listed in Table S1.

The small GTPase RAB8 functions in ciliary membrane extension during later steps in ciliogenesis.^9^ We previously showed that IFT20 is reduced at the centrosome in patient-derived trisomy 21 fibroblasts.^19^ IFT20 is recruited to the mother centriole upstream of RAB8 and is required for RAB8 targeting to the ciliary membrane.^9,21^ We thus examined RAB8 localization to the mother centriole during the ciliogenesis time course to determine if later trafficking pathways are also disrupted. By 8 hours post serum starvation, T21 and Q21 cells showed decreased RAB8 accumulation at the mother centriole compared to D21, and this decrease persisted throughout the time course (Figures 4C-D, S4D). Given that RAB8 associates with Golgi and post-Golgi membranes in the context of ciliogenesis,^29,30^ one potential explanation for decreased RAB8 at the centrosome is defective trafficking from the Golgi. However, we did not observe changes in RAB8 intensity at the Golgi with increasing ploidy, suggesting RAB8 leaves the Golgi without incident (Figures S4E-F). Thus, RAB8 trafficking defects are specific to the centrosomal region and are not generally disrupted intracellularly in T21 and Q21 cells.

Given the observed defects in trafficked proteins required for ciliogenesis reaching the centrosome and the importance of microtubules in trafficking pathways, we wondered if general membrane trafficking pathways were disrupted with increasing HSA21 ploidy. *Cis*- and *trans*-Golgi networks, early endosomes, and lysosomes did not exhibit changes in morphology in T21 and Q21 cells when compared to D21 cells (Figures S4G-Q). Moreover, whereas mild intensity differences were observed in some of these organelles, they did not follow a consistent trend with increasing ploidy (Figures S4G-Q). Thus, gross disruptions of the endolysosomal trafficking pathways are not apparent and trafficking defects associated with trisomy 21 are focused at and around the centrosome with increasing HSA21 ploidy. Collectively, this suggests that ciliogenesis defects in trisomy 21 result from early PCNT-induced crowding around the centrosome that captures preciliary vesicle components thereby preventing mother centriole uncapping and RAB8-axoneme extension.

### Decreased transition zone protein localization in ciliated trisomy 21 cells

Despite early defects in ciliogenesis, elevated PCNT from increasing HSA21 ploidy does not abolish cilia. Approximately 40% of T21 and 20% of Q21 cells formed a primary cilium by 24 hours post serum depletion (Figure 1E). We thus wondered whether trisomy 21 induced a purely kinetic block in cilia formation or if defects persisted after ciliation. To function as a signaling compartment, primary cilia are exposed to the external cellular environment to send and receive signals. In RPE1 cells, cilia assemble inside the cell and then fuse with the cell membrane to become extracellular signaling organelles.^31^ This requires remodeling of the plasma membrane and cytoskeletal networks. Given the observed centrosomal trafficking defects, we tested whether cilia in trisomy 21 cells were exposed to the external environment using the IN/OUT assay.^32^ No difference was observed in the number of cilia outside versus inside the cell in D21 and T21 cells (Figure S5A-B), suggesting that trisomy 21 does not cause cilia to be retained inside cells.

A second requirement for cilia function in signaling is the creation of a diffusion barrier called the transition zone. The transition zone, at the base of the cilium, is composed of proteins that regulate entry and exit of molecules to and from the cilium. Regulation of diffusion through the transition zone is essential for proper ciliary signaling, and many genes mutated in ciliopathies encode transition zone proteins.^33,34^ The levels of transition zone proteins were analyzed in D21 and T21 cells. Q21 cells were eliminated because of their low ciliation frequency. Consistent with defects in the primary cilia transition zone, the core transition zone protein CEP290^35,36^ was decreased in T21 cells compared to D21 cells (Figures 5A, yellow arrows, and 5B). In addition to the transition zone, CEP290 also localizes to centriolar satellites^13^, and we observed increased CEP290 at the pericentrosomal but not centrosomal region in T21 cells relative to D21s (Figures 5A, 5C, S5C). Moreover, CEP290 colocalizes with PCNT in this pericentrosomal region (Figure 5A, cyan arrows), suggesting that decreased CEP290 at the transition zone may result from pericentrosomal crowding. In addition to CEP290, RPGRIP1L, and two outer transition zone proteins, NPHP4 and TMEM67,^37^ were also decreased at the transition zone in T21 compared to D21 cells (Figures 5D-I). Collectively, trisomy 21 cells show transition zone defects suggesting that trafficking problems to the centrosome are not specific to building a cilium but persist even in ciliated cells. This may explain the signaling defects found previously in cells with elevated PCNT.^19^

**Figure 5.**
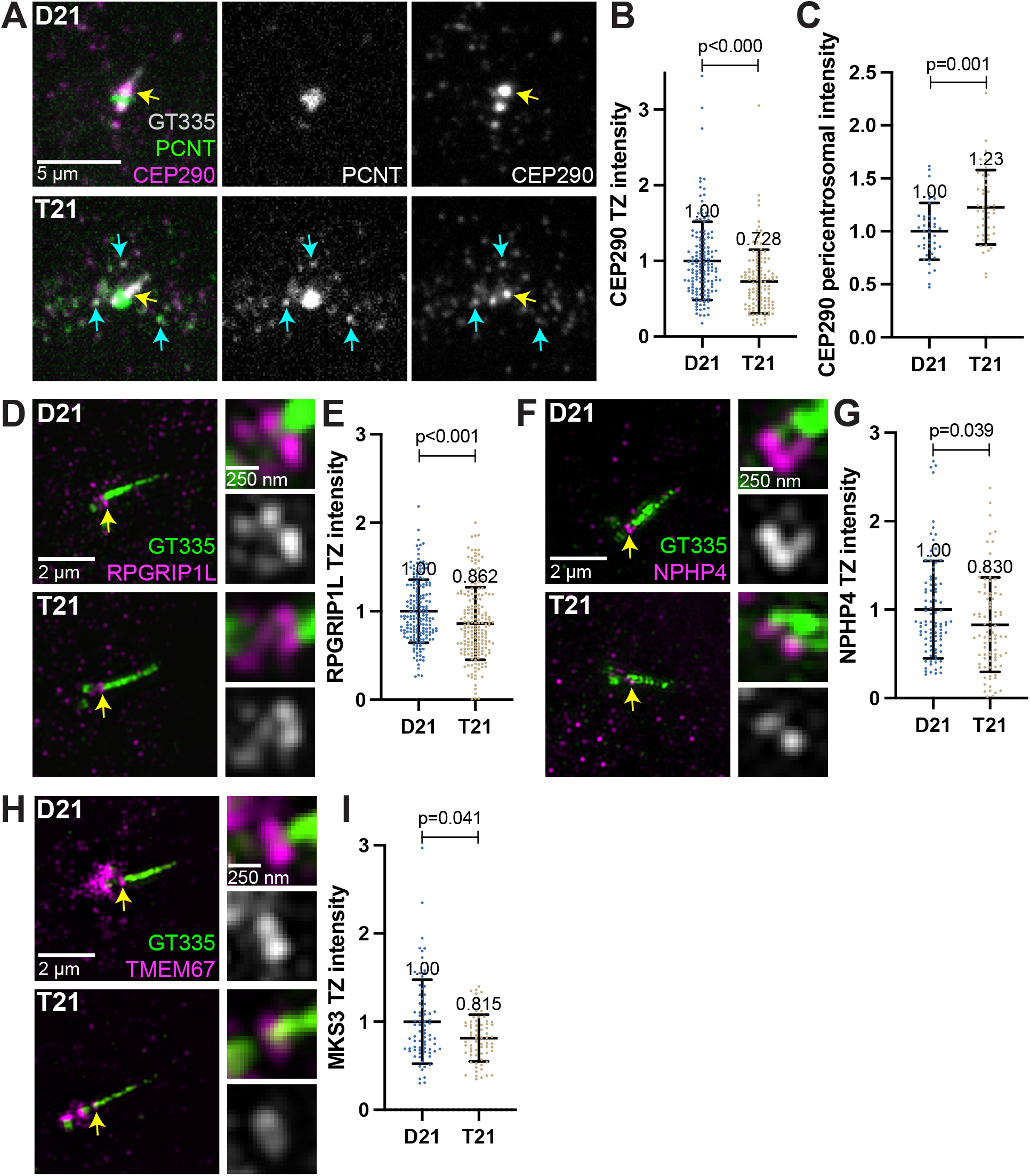
Decreased transition zone protein localization in ciliated trisomy 21 cells. (A) Representative confocal images of RPE1 D21 and T21 cells grown on coverslips and serum depleted for 24 h. Cells were stained with GT335, PCNT, and the transition zone protein CEP290. Yellow arrows point to the CEP290 transition zone population and cyan arrows point to CEP290 satellites that colocalize with PCNT. (B) Quantitation of CEP290 transition zone intensity. (C) Quantitation of pericentrosomal CEP290 intensity in 1.2-5 μm region around centrosome. (D, F, H) Representative SIM images of RPE1 D21 and T21 cells grown on coverslips and serum depleted for 24 h. Cells were stained with GT335 and the transition zone proteins RPGRIP1L (D), NPHP4 (F), or TMEM67 (H). (E, G, I) Quantitation of indicated transition zone protein intensities from confocal images. All values were normalized to the D21 average, and graphs show mean ± SD. Mean intensity values are indicated on graphs. N’s and statistical tests are listed in Table S1.

### Shh signaling is defective in primary mouse fibroblasts with elevated PCNT

The barrier function of the transition zone is critical for proper ciliary signaling.^34,38^ Because transition zone defects were found in trisomy 21 cells, we next asked whether trisomy 21 negatively affects primary cilia dependent signaling. Shh signaling is the best understood cilia-dependent signaling pathway and is important for developmental events that are impacted in DS. Moreover, Shh signaling was previously found to be disrupted in a DS mouse model.^39^ Shh signaling is commonly studied in primary mouse embryonic fibroblasts (MEFs). HSA21 maps to syntenic regions of three mouse chromosomes: MMU10, MMU16, and MMU17.^40^ We analyzed three mouse models harboring segmental genomic duplications of these syntenic regions on MMU10, MMU16, or MMU17 called Dp10, Dp16, or Dp17, respectively.^41^ The murine *Pcnt* gene is located on MMU10, therefore only the Dp10 model contains an extra copy of the *Pcnt* gene, although other cilia and centrosome-related genes can be found on MMU16 and MMU17 (Figure 6A).^19,42^

**Figure 6.**
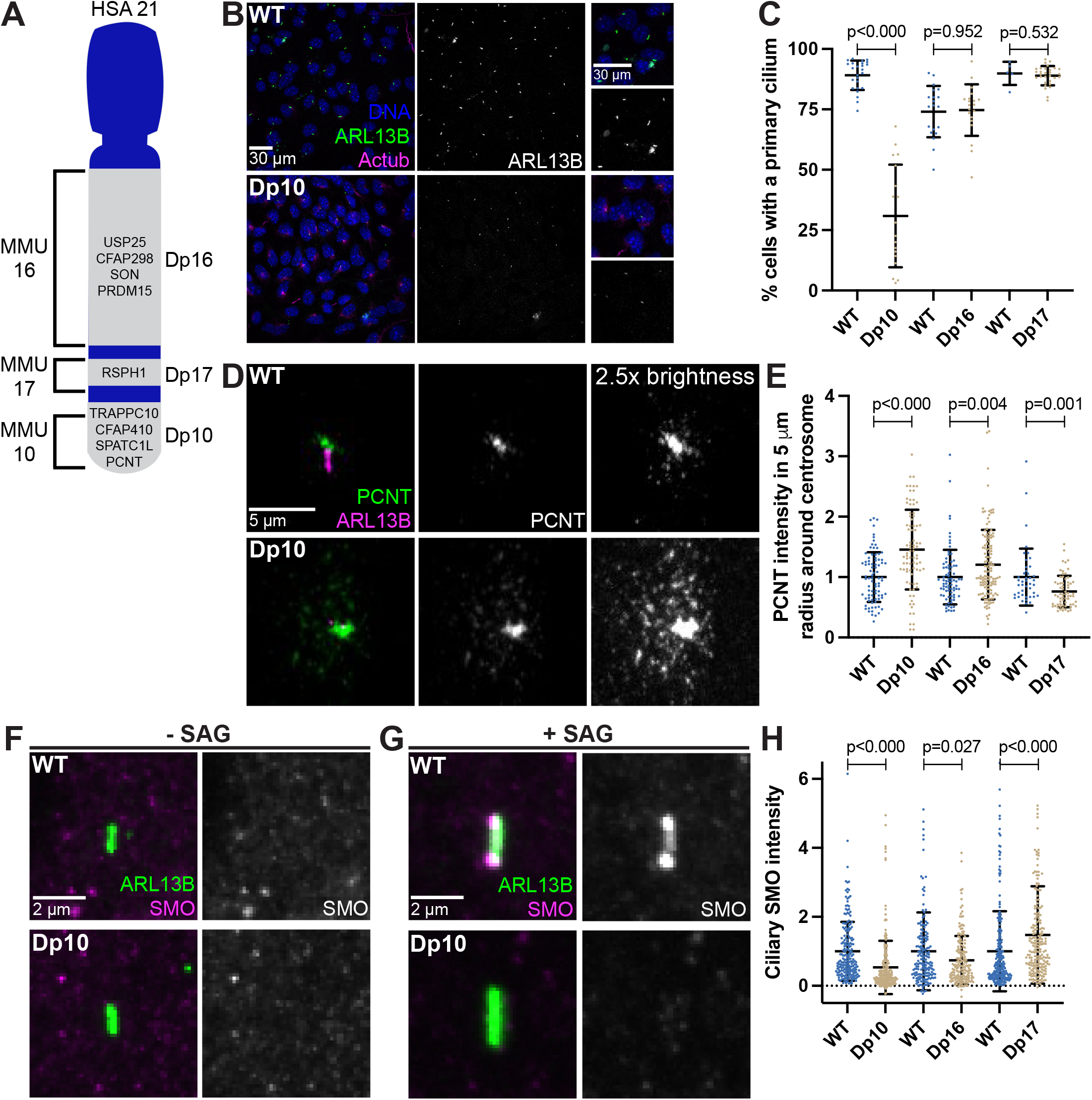
Shh signaling is defective in primary mouse fibroblasts with elevated PCNT. (A) Cartoon model depicting mouse syntenic regions with HSA21 and corresponding Dp10, Dp16, and Dp17 mouse models. PCNT is located on MMU10. Other cilia and centrosome proteins are also listed. For all following experiments, WT and Dp primary MEFs were isolated from E12.5 dams, grown on coverslips, and serum depleted for 24 h. (B) Representative confocal images of WT and Dp10 MEFs. Cells were stained with Hoechst, the ciliary marker ARL13B, and Actub. (C) Quantitation of the number of cells with a primary cilium in WT and Dp MEFs. (D) Representative confocal images of WT and Dp10 MEFs. Cells were stained with PCNT and ARL13B. (C) Quantitation of PCNT intensities in a 5 μm radial circle around the centrosome in WT and Dp MEFs. Values were normalized to the WT average. (F, G) Representative confocal images of WT and Dp10 MEFs untreated (F) or treated with 100 nM SAG for the last 4 h of serum depletion (G). Cells were stained with SMO and ARL13B. (H) Quantitation of ciliary SMO levels in SAG-treated cells for WT and Dp MEFs. Graphs show mean ± SD. N’s and statistical tests are listed in Table S1.

We first asked whether MEFs isolated from Dp10, Dp16, or Dp17 embryos showed ciliation defects. Strikingly, only the Dp10 MEFs with elevated PCNT showed decreased primary cilia frequency when compared to wild-type littermates (Figures 6B-E). Cilia frequency in Dp16 and Dp17 MEFs was identical to wild-type (Figures 6C, S6A-B). Dp10 mice contain approximately 41 duplicated HSA21 gene orthologs,^41^ and we previously tested the other cilia and centrosome human orthologs on MMU10 and found no changes in ciliation.^19^ Thus, decreased ciliation is specific to elevated PCNT in primary MEFs from the Dp10 mouse model of DS.

Upon induction of the Shh pathway, the transmembrane protein Smoothened (SMO) translocates into the cilium and generates signals that induce nuclear GLI localization and downstream transcriptional response of hedgehog target genes important for mitogenic activity and developmental processes such as patterning and limb development.^4,43^ Because Shh signaling is disrupted in a DS mouse model,^39^ and we previously found decreased GLI expression in DS-derived human fibroblasts,^19^ we next examined Shh signaling in MEFs where we can distinguish the contributions of different regions of HSA21. We compared SMO intensity in the cilium with and without induction of Shh signaling using the SMO agonist SAG. Without SAG, ciliary SMO levels were undetectable in both wild-type and Dp10 MEFs (Figures 6F and 6H). However, upon SAG treatment, SMO robustly accumulated in the cilium in wild-type cells, whereas ciliary SMO remained low in Dp10 cells (Figures 6G-H). Moreover, ciliary SMO levels in Dp16 and Dp17 cell lines after SAG treatment were the opposite of PCNT levels, as Dp16

MEFs had slightly elevated PCNT levels and slightly decreased ciliary SMO while Dp17 MEFs had slightly decreased PCNT levels and slightly increased ciliary SMO (Figures 6D-E, 6H, S6C-D). Together, decreased ciliation frequency and ciliary SMO in Dp10 MEFs is consistent with persistent trafficking defects from pericentrosomal crowding in ciliated trisomy 21 cells. Moreover, PCNT levels anticorrelate with ciliary SMO in Dp16 and Dp17 MEFs.

Interestingly, one Dp10 MEF line (Dp10-2) did not show decreased ciliation compared to wild-type controls (Figures S6E-F). Importantly, this line also did not exhibit elevated PCNT levels or defects in ciliary SMO localization upon SAG treatment (Figures S6G-H). It is unclear whether this Dp10-2 line lost the chromosome duplication containing PCNT or whether cells compensated at the molecular level. Regardless, results from this line reinforce the conclusion that cilia and signaling defects result from elevated PCNT levels. In summary, PCNT and ciliary SMO levels anticorrelate and are disrupted in trisomy 21.

### Elevated PCNT in a DS mouse model results in decreased primary cilia and cerebellar dysmorphology

Primary cilia are ubiquitous and essential signaling organelles. They are particularly important during brain development and are predicted to be disrupted in DS.^4,44^ Individuals with DS commonly exhibit cerebellar hypoplasia,^1^ and delayed cerebellar development has been observed in the Ts65Dn mouse model of DS, which harbors a duplication of MMU16 genes similar to that of the Dp16 model along with a duplication of genes from MMU17 that are not syntenic to human HSA21.^39,40^ During cerebellar development, neuronal precursor cells in the external granular layer respond to Shh from Purkinje cells.^45,46^ Shh induces mitogenic activity, and the amplified cells ultimately migrate to the internal granule layer where they become mature neurons.^46^ Primary cilia are required for neuronal precursor cell amplification and defects in Shh signaling reduce proliferation and disrupt cerebellar development.^47^ Yet, whether T21 and elevated PCNT disrupt primary cilia during brain development *in vivo* remains unknown. We therefore examined ciliation in the external granular layer of postnatal day 4 (P4) pups. Consistent with our findings in Dp10 MEFs, cerebellar neuronal precursors in Dp10 animals had fewer primary cilia compared to wild-type littermates (Figures 7A, 7D). Moreover, Dp16 and Dp17 animals showed no change in ciliation frequency (Figures 7B-D). These data are consistent with our analyses in MEFs suggesting that cilia defects occur in animals with elevated PCNT.

**Figure 7.**
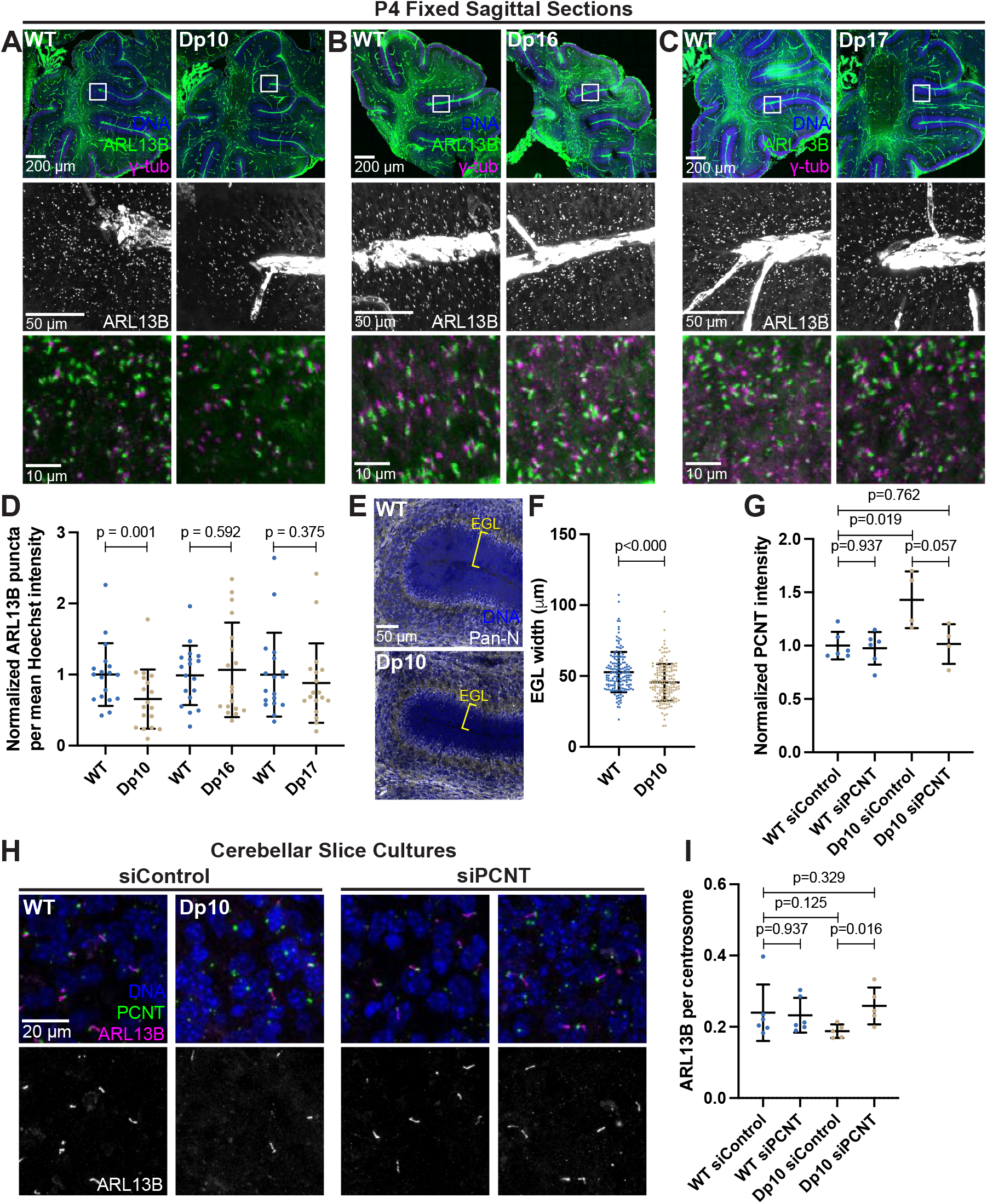
Elevated PCNT in a DS mouse model results in decreased primary cilia and cerebellar dysmorphology. (A-C) Representative tiled confocal images of the cerebellum from P4 wild-type (WT) and Dp10 (A), Dp16 (B), and Dp17 (C) animals. Brain sections were stained with Hoechst, ARL13B, and γ-tubulin. Insets show progressively zoomed in regions corresponding to the same folia in each animal. (D) Quantitation of primary cilia frequency in WT and Dp animals normalized to WT. (E) Representative tiled confocal images of WT and Dp10 P4 animals corresponding to the same cerebellar folia in each animal. Brain sections were stained with Hoechst and a Pan-neuronal marker. Yellow bracket denotes external granular layer. (F) Quantitation of the external granular layer width in WT and Dp10 animals. (G) Quantitation of primary cilia frequency in WT and Dp10 cerebellar slice cultures treated with control or PCNT siRNA. Values were normalized to WT siControl averages. (H) Representative confocal images of WT and Dp10 cerebellar slice cultures isolated from P4 pups and treated with control or PCNT siRNA for 48 h in serum free media. Slices were stained with Hoechst, ARL13B, PCNT. Yellow box denotes insets. (I) Quantitation of PCNT intensity in WT and Dp10 cerebellar slice cultures treated with control or PCNT siRNA. Values were normalized to WT siControl averages.

Reduced Shh signaling correlates with morphological changes in the cerebellum, such as decreased width of the external granular layer.^48,49^ We thus measured the width of the external granular layer and found decreased widths in Dp10 animals compared to wild-type littermates (Figure 7E-F). Decreased width could result from decreased cell proliferation, so we next examined Ki67 staining as a marker for cell proliferation. While we observed no changes in the number of Ki67-positive cells in the external granular layer of Dp10 and wild-type animals at P4 (Figure S7A-B), we cannot rule out changes in cell proliferation at earlier developmental time points. The source of Shh for neuronal precursors comes from Purkinje cells,^46^ so we next examined the Purkinje cell layer. No loss of Purkinje cells was noted between Dp10 and wild-type siblings (Figure S7C). Finally, given that elevated PCNT alters microtubule networks in cultured RPE1 cells and that PCNT mutations produce severe neuronal defects including disrupted neuronal migration,^18,50,51^ we visualized doublecortin (DCX), a microtubule binding protein that functions in neuronal migration.^52,53^ Interestingly, Dp10 animals showed decreased DCX-labeled cellular protrusions compared to wild-type (Figure S7C). While not conclusive, this suggests that elevated PCNT alters non-centrosomal microtubules such as those required for neuronal outgrowth.^54^ Taken together, elevated PCNT results in decreased ciliation in the external granular layer of the cerebellum and these cerebella demonstrated morphology changes consistent with decreased Shh signaling and developmental defects.

To determine if decreased ciliation *in vivo* was a direct result of elevated PCNT levels, we cultured cerebellar slices from P4 pups and treated slices with either control or PCNT siRNA. Strikingly, reducing PCNT levels in Dp10 slice cultures rescued ciliation back to control levels (Figure 7G-I). Taken together, this suggests that elevated PCNT from trisomy 21 is sufficient to induce ciliation defects during cerebellar development *in vivo*.

Our data from T21 and Q21 RPE1 cells suggested that pericentrosomal crowding delayed ciliogenesis; however, cells arrested in G1 for a prolonged period eventually ciliated albeit at slightly lower frequencies than D21 cells (Figure 1E). Moreover, our previous work in human DS-derived fibroblasts showed the strongest ciliation defects in cycling cells.^19^ We thus tested whether ciliation defects *in vivo* were dependent on cell cycle state by analyzing cells in the inner granular layer of the cerebellum which are generally post-mitotic. Primary ciliation of inner granule layer cells was unchanged between wild-type and Dp10 animals (Figure S7D). This provides further evidence that trisomy 21 alters but does not abolish ciliogenesis *in vivo,* with the major impact on migrating precursor cells

## Discussion

### Trisomy 21 alters trafficking flux of early ciliogenesis molecules delaying ciliogenesis

Here, we use a time course to establish the dynamic changes to PCNT trafficking puncta and microtubules upon induction of ciliogenesis through media serum depletion. In control euploid cells, the largest changes to PCNT and microtubule networks around the centrosome occur early, within 2 h after induction of ciliogenesis. Moreover, we observe a slow and a fast phase to ciliation in a population of cells, with the fast phase occurring within the first 8 h and the slow phase from 8-48 h. In T21 and Q21 cells, PCNT and microtubule levels increase more dramatically at the onset of ciliogenesis. This is accompanied by a decrease in the effectiveness of the fast phase of ciliogenesis, while the slow phase from 8 to 48 h is largely the same as D21 cells. These data reveal that in trisomy 21, PCNT-induced pericentrosomal crowding occurs immediately upon induction of ciliogenesis and delays but does not abolish ciliation.

Several distinct trafficking pathways required for building a primary cilium are prevented from efficient recruitment to the site of assembly in the altered microtubule and PCNT landscape caused by increased HSA21 dosage. MYOVA, one of the earliest proteins to localize to the mother centriole, does not traffic normally but rather participates in pericentrosomal crowding in T21 and Q21 cells. Mother centriole uncapping is also disrupted, perhaps through defects in trafficking the ubiquitin or autophagosome machinery required for CP110 and CEP97 removal and degradation, although this remains to be tested. In addition to early defects in ciliogenesis, pericentrosomal crowding affects later steps such as RAB8-mediated ciliary membrane growth and IFT20 centrosomal localization.^9,19^ Together, our results show HSA21 dosage dependent disruption in trafficking pathways important to the earliest observable stages of ciliogenesis and cilia function, as molecules required for building and maintaining a cilium get hung up in PCNT-induced crowding around the centrosome.

Consistent with a crowding model, several lines of evidence support the idea that trisomy 21 does not result in a complete block to ciliogenesis, but rather a delay in the process. First, with sustained G1 arrest, T21 and Q21 cells increase their ciliation, and these cells appear to compensate for crowding such that the difference between PCNT and microtubule densities at and around the centrosome is reduced between D21, T21, and Q21 cell lines. Second, a similar phenomenon is observed with crowding cargo, as the amount of pericentrosomal MYOVA decreases over time and MYOVA eventually accumulates at mother centrioles in T21 and Q21 cells. Third, trisomy 21 cells demonstrate defective signaling even after a cell builds a primary cilium. This is consistent with previous work showing reduced GLI expression in human trisomy 21 cells and decreased ciliary SMO trafficking in cells overexpressing PCNT.^19^ Fourth, crowding has the most severe consequences for molecules that are dynamically trafficking to and from the centrosome, whereas longer lived structures such as the core centriole and appendages^55^ are unaffected by elevated PCNT. Finally, *in vivo*, only cycling neuronal precursors of the external granular layer have decreased ciliation, whereas ciliation in post-mitotic cells of the inner granular layer remains unaffected with trisomy 21. Collectively, we find that PCNT-induced pericentrosomal crowding disrupts the dynamic flux of molecules to and from the centrosome thereby disturbing the coordination and timing required for proper ciliogenesis and signaling.

### Cerebellar phenotypes in animals with elevated PCNT

While the timing of ciliogenesis is not vital in cultured cells, the timing of ciliogenesis and cilia-dependent signaling is critical during *in vivo* development where cilia send and receive input from neighboring cells to coordinate proper tissue development.^44^ In line with our cultured cell results, we do not observe a complete loss in cilia and signaling in primary MEFs or cerebellar neuronal precursor cells with elevated PCNT but do observe consistent decreases in ciliation. Moreover, the organization of cerebellar layers with a thinner external granular layer is altered but not eliminated. While we might expect to see decreased cell proliferation in cells in the external granular layer, Ki67 staining, a marker for proliferating cells, appears normal in Dp10 animals compared to wild-type littermates. This result could occur for several reasons. First, our analysis was conducted only in P4 pups, where developmental delays are not yet obvious in Dp10 animals. An analysis of cell proliferation at earlier or later timepoints might show changes to the number of cycling cells. Second, because elevated PCNT does not result in a complete loss of cilia, perhaps enough of the cell population is ciliated to receive the Shh mitogenic signal and proliferate at this timepoint. Cerebellar development is a highly integrated process where a trisomy 21-induced delay in ciliogenesis could alter the coordination of cellular processes such that some cells never fully catch up. Because inputs are not completely lost, this cell-to-cell variability could account for the spectrum of phenotypes observed in individuals with DS.

Most trisomy 21 animal studies have been performed in the Ts65Dn mouse model.^40^ While these mice show some degree of DS-like phenotypes, they are only trisomic for about half of the HSA21 orthologs and contain an additional amplification of genes not found on HSA21.^40,56^ The Dp mouse models are a more refined genetic system to determine the phenotypic contributions from different regions of HSA21. Dp10 mice contain approximately 41 duplicated HSA21 gene orthologs including PCNT while Dp16 and Dp17 mice have 115 and 19 duplicated gene orthologs, respectively, and exclude PCNT.^41^ Our experiments in primary MEFs and cerebellar slices largely attribute cilia defects to elevated PCNT levels in Dp10 animals. However, there are other known cilia and centrosome genes found on HSA21, and potentially non-coding regions that may also contribute to phenotypic consequences of trisomy 21. For example, MMU16 contains the splicing factor SON, which is known to splice PCNT mRNA and alters PCNT levels and distributions.^57,58^ Indeed, while we do not find changes in primary cilia frequency in Dp16 animals, we do observe moderately decreased ciliary SMO and moderately increased PCNT levels. Interestingly, Dp16 mice have defects in the motile cilia lining the ependymal cells of the brain (Figure S7E-G) suggesting that elevated gene dosage can have different phenotypic consequences in different tissues. In support of this, animals with individual chromosomal duplications, Dp10, Dp16, or Dp17,^41^ do not show as severe phenotypic consequences as combined Dp10; Dp16; Dp17 animals.^59^ Therefore, while elevated PCNT disrupts ciliogenesis and signaling, there are likely other contributions from additional genes on HSA21 that add to the spectrum of phenotypes observed in individuals with DS.

In contrast to single gene disruption studies, here we show that ciliary defects arise from elevated protein expression, as is often the case in chromosomal aberrations. Indeed, cilia defects have been observed with increased copy number of a nuclear pore protein.^60^ While changes in some protein levels may be tolerated, we demonstrate that even a modest increase in PCNT protein (1.5 - 2-fold) is deleterious to cilia formation and function in a dose dependent manner. Elevated PCNT aggregates and reorganizes microtubule networks, inducing pericentrosomal crowding that disrupts cargo transport required for early steps in ciliogenesis and persists in ciliated trisomy cells. Because tight control of microtubule organization is essential for many cell types and processes including neuron outgrowth, cell migration, immune synapse formation, and cell polarity, these functions need further study to determine whether elevated PCNT from trisomy 21 and changes in microtubule topologies alter these cell types and functions.

## Acknowledgements

We thank Drs. Andrew Lane and David Pellman for RPE1 D21, T21, and Q21 cell lines, Dr. Chris Westlake for EHD1 plasmid, Dr. Rajat Rohatgi for SMO antibody, Dr. Tamara Caspary for advice on isolating primary MEFs, and Dr. Derek Toomre for pH-Smoothened plasmid. We are grateful to Drs. Carolyn Ott, Santos Franco, and the Pearson lab for helpful discussions. Electron microscopy was done at the University of Colorado, Boulder EM Services Core Facility in the MCDB Department with Garry Morgan providing specimen preparation. This research was funded by NIH R01GM138415 and R35GM140813 to CGP, NSF Graduate Research Fellowship DGE-1553798, NIH INCLUDE T32 supplement GM008730, and Blumenthal Fellowship to CEJ, and NIH R01DK064380 to RP. CEJ, JME, KDS, and CGP are members of the Linda Crnic Institute for Down syndrome.

## Author Contributions

CEJ, BLM, ETO, KSG, CHL, and VO performed experiments and analyzed data. CEJ and CGP conceived the project and wrote the manuscript. WM, LR, JME, KDS, WBM, and RP provided reagents, animal models, and expertise.

## Abbreviations

HSA21: human chromosome 21
MMU: mouse chromosome
PCNT: pericentrin
DS: Down syndrome
Shh: Sonic hedgehog
MEF: mouse embryonic fibroblast
RPE1: retinal pigmented epithelia

## Materials and Methods

### Cell lines

Disomy 21, Trisomy 21, and Tetrasomy 21 (D21, T21, Q21) hTERT-immortalized retinal pigment epithelial (RPE1) cells were generated by and Drs. Andrew Lane and David Pellman at the Dana-Farber Cancer Institute.^61^ Cells were grown in DMEM:F12 (SH30023; Cytiva) supplemented with 10% fetal bovine serum (FBS, Peak Serum; PS-FB2) and 1% Penicillin/Streptomycin at 37°C and 5% CO_2_. Cells were passaged 1:5 at ~80-90% confluency with 0.25% Trypsin (15090-046; Gibco). Cells were routinely screened for mycoplasma. Primary MEFs were generated as described in Mariani et al.^62^ In brief, embryonic day 12.5 embryos were dissected from pregnant females. The head and internal organs were removed, and the remaining tissue was dissociated by passage through a needle. The head was used for genotyping. Dissociated cells were cultured in DMEM:F12 with 10% fetal bovine serum and 1% Penicillin/Streptomycin for no more than 3 passages.

### Immunofluorescence

Cells were plated on collage-coated coverslips and grown in full media until 80-90% confluent. To induce ciliogenesis, cells were washed once with 1x PBS and then grown in serum depleted media (DMEM:F12 with 0.5% FBS) for indicated hours. For time course experiments, cells were fixed according to Waterman-Storer and Salmon^63^ following 0, 2, 4, 8, 24, or 48 hours post serum depletion. Briefly, cells were pre-permeabilized for 5 minutes in 0.5% TritonX-100 in PHEM (60mM Pipes, 25mM Hepes, 10mM EGTA, 2mM MgCl_2_, 6.9 pH). Cells were then fixed with 4% paraformaldehyde/0.5% glutaraldehyde for 20 minutes and quenched with 0.1% sodium borohydride. Following quenching, cells were washed four times in 0.1% TritonX-100 and stored in PHEM at 4°C until immunostaining.

For all other staining experiments, cells were fixed with 4% paraformaldehyde for 20 minutes at room temperature or with 100% ice cold Methanol for 10 minutes at −20°C. Cells were blocked for 1-2 hours in block buffer (PBS, 0.1% Triton X-100, 10% normal donkey serum). Primary antibodies were diluted in block buffer and incubated overnight at room temperature. Cells were washed with PBSTx before adding secondary antibodies for 1-2 hours at room temperature. Cells were washed again before mounting in VectaShield and sealing with nail polish or mounting in Prolong Gold for SIM experiments. Coverslips for all experiments were #1.5. Primary and secondary antibodies are listed in the Key Resources Table.

For SAG treatment, MEFs were grown on coverslips to 80-90% confluency, moved to serum depleted media for 20 hours, treated with 100 nM SAG in serum depleted media for 4 hours, fixed with 4% PFA and stained as described above.

To costain PCNT with other rabbit primary antibodies, PCNT was conjugated to Alexa Fluor 488 (Antibody labeling kit: Invitrogen A20181).

### Fluorescence microscopy

Confocal images were acquired using either a Nikon Eclipse Ti 2 inverted A1 Confocal and Nikon Elements software or Nikon Eclipse Ti inverted microscope stand equipped with a 100x Plan Apochromat objective (NA 1.45), Andor iXon X3 camera, CSU-X1 (Yokogawa) spinning disk, and Slidebook 6 software. Super resolution imaging was performed on a Nikon 3D SIM system (Ti 2 Eclipse) with a 100x TIRF objective (NA 1.45). Images were captured with a complementary metal-oxide semiconductor camera (Orca-Flash 4.0; Hamamatsu). Raw images were reconstructed using the Nikon Elements image stack reconstruction algorithm. Images were processed in FIJI ^64^. Figures were made in Adobe Illustrator. A minimum of three biological replicates were performed for each experiment unless otherwise noted. All images presented in figures are max projections.

### IN/OUT cilia assay

RPE1 D21, T21, and Q21 cells stably expressing pHluorin-Smoothened (pLVX-pHluorin-Smoothened vector was a gift from Dr. Derek Toomre) were plated on collagen-coated coverslips, fixed and stained as described in^32^. Briefly, culture media was removed, cells were gently washed in PBS, and then fixed in 4% PFA in PBS for 10 minutes. Cells were then blocked in 5% normal donkey serum in PBS for 30 minutes followed by incubation with anti-GFP primary antibody solution for 1 hour. Cells were fixed again for 10 minutes, permeabilized with 0.1% Triton-X, and incubated in another primary antibody solution containing anti-Actub for 1 hour. After gentle washing, cells were incubated with secondary antibodies and Hoechst, followed by washing and mounting on slides.

### RNAi

Human PCNT siRNA (Smart Pool) (M-012172-01-0005; Dharmacon) was transfected into RPE1 cells with Lipofectamine RNAi MAX (13778100; ThermoFisher Scientific) according to the manufacturer’s protocol. Mission siRNA universal negative control #1 was used for all negative controls (SIC001-1NMOL; Sigma). All siRNAs were used at a final concentration of 25 nM. Cells were treated with siRNA in serum depleted media for 24 hours before fixation and subsequent immunostaining steps.

### Generation of lentiviral stable cell lines

RPE1 D21, T21, and Q21 lentiviral stable cell lines were generated by transfecting HEK293T cells with lentiviral GFP-EHD1 or pH-Smoothened constructs and lentivirus packaging plasmids using Lipofectamine 2000 (11668-027; Invitrogen). HEK293T media containing virus was collected and added to target cells in the presence of 2mg/mL polybrene. Transduced cells were selected using 10mg/mL puromycin for three days. To induce GFP-EHD1 expression, 0.125 mg/mL doxycycline was added to serum depleted media for 24 hours before fixation.

### Animal Ethics Statement

All procedures involving mice were approved by the Institutional Animal Care and Use Committee (IACUC) at the University of Colorado Anschutz Medical Campus and were performed in accordance with National Institute Guidelines for the care and use of animals in research.

### Mouse models of Down syndrome

Mouse models were obtained from the NICHD funded Cytogenetic and Down Syndrome Models Resource Jackson Laboratory. These include B6.129S7-Dp(16Lipi-Zbtb21)1Yey/J (Dp16; stock# 013530), B6;129-Dp(10Prmt2-Pdxk)2Yey/J (Dp10; stock# 013529), and B6;129-Dp(17Abcg1-Rrp1b)3Yey/J (Dp17; stock# 013531). Strains were maintained on a standard chow diet and a 14-hour light/10-hour dark cycle. Littermate controls were used.

### Cerebellar Slice Cultures

Cerebellums were dissected out of postnatal day 4 pups in ice cold Hank’s media and sliced 300 μm thick on a tissue chopper.^65^ Slices were transferred to MilliCell filter inserts (Millipore PICM03050) only taking intact slices with at least 5 folia. Excess Hank’s was removed from filters and replaced with serum free Neurobasal media supplemented with 1% N2, 0.5% Penicillin/Streptomycin, 0.25% L-glutamine, and 10 mM HEPES. Cerebellar slices were cultured at 37°C and 5% CO_2_ and media was changed 8 hours after plating. Slices were grown for 48 hours before addition of 500 μm control or PCNT siRNA (D-001910-10-05 Accell Non-targeting Pool; Accell Mouse Pcnt 18541 siRNA—SMARTpool) and then cultured for an additional 48 hours before fixation with 4% PFA and staining as described above.

### Mouse brain immunohistochemistry

Experiments were blinded to gender. P4 pups were anesthetized with Isoflurane before decapitation. Tail clips were used for genotyping. Brains were dissected out and fixed in 4% PFA in PBS overnight at 4°C. Brains were then moved to 30% sucrose in PBS for 1-2 days at 4°C. Fixed brains were sliced through the midline; each half was embedded in OCT, frozen on dry ice for ~15 minutes, then stored at −80°C until ready to section. To section, blocks were mounted in OCT and 20 μm sagittal sections were cut using a Leica CM 1950 cryostat microtome. Sections were placed on FisherBrand charged slides (Cat # 12-550-15) and stained on slides as described above.

### Genotyping

Tissue from Dp10 animals or MEFs were lysed by placing tail clips or heads in Gitschiers Buffer (67 mM Tris pH 8.8, 0.166 mM Ammonium sulfate, 6.7 mM magnesium chloride, 0.005% TritonX, 0.1 mg/ml Proteinase K in water) for 2 hours at 55°C then 95°C for 10 minutes. Tissue from Dp16 or Dp17 animals or MEFs were lysed by placing tail clips or heads in 50 mM sodium hydroxide and heating at 98°C for 2 hours before neutralizing with 1M Tris pH 8 (1:10). Lysed tissue was used for PCR and the resulting banding pattern from PCR was used to determine genotype. Genotyping primers are as follows: Dp10For: GGCGAACGTGGCGAGAAA; Dp10Rev: CCTGCTGCCAAGCCATCAG; Dp16For: CTGCCAGCC ACTCTAGCTCT; Dp16Rev: AATTTCTGTGGGGCAAAATG; Dp17For: GGAGCCAGGGCTGATGGT; Dp17Rev: CAACGCGGCCTTTTTACG. Primers for Cux2 were used as controls. Cux2For: GGGACATCACCCACCGGTAATCTC; Cux2Rev: GACCACTGAGTCTGGCAACACG.

### Image Analysis

All intensity analysis was performed on max projected images unless otherwise noted. Radial fluorescence intensity was measured using the Radial Profile Extended ImageJ plugin. Briefly, this plugin plots average fluorescence intensity as a function of distance from a user defined centroid. 10 μm diameter ROIs were centered over the brightest pixel of a Gaussian blurred maximum intensity projection of the centrosome defined by PCNT fluorescence. In cells with GT335-labeled centrioles, the centroid was manually centered between the two centrioles. Background was determined per field of view by the mean intensity of an extracellular ROI. Background subtracted radial intensities were summed and normalized to the D21 average. Centrosomal fluorescence intensity was defined as the sum of radial intensities falling within 0.0-1.2 μm from the centroid of analysis. Pericentrosomal fluorescence intensity was defined as the sum of radial intensities falling within 1.2-5.0 μm from the centroid of analysis.

Whole cell intensities were calculated using the integrated density within cells outlined in ImageJ. Ciliation frequency was measured by counting either DM1a (Figure 1) or ARL13B (Figures 6) and dividing by the number of nuclei per field of view. For ciliation frequency in cultured cerebellar slices, the number of ARL13B cilia was divided by the number of centrosomes per field of view. For ciliation frequency in fixed cerebellar slices, the number of ARL13B cilia was divided by the mean Hoechst intensity per field of view because individual nuclei could not be resolved in the thick slices.

For PCNT siRNA experiments, radial MYOVA and PCNT fluorescence intensity was calculated using the same methods as above. For MYOVA radial distribution, all values were divided by the maximum intensity value per cell and normalized to D21 values. Correlation analyses were performed by plotting 10 μm diameter PCNT intensities normalized within a cell line to either centrosomal or pericentrosomal normalized MYOVA intensities. R values and significance were calculated by running a correlation analysis in GraphPad. PCNT levels were analyzed with a 10 μm diameter circle centered on the centrosome to measure the integrated density. Background was determined with an extracellular ROI. Background subtracted values were normalized to the D21 average.

GM130 and Golgin97 intensities were measured by drawing an ROI around the GM130 or Golgin97 signal and measuring the integrated density within this ROI. Background was determined by an extracellular ROI. Background subtracted values were normalized to the D21 average. To quantify early endosome and lysosome levels, integrated density was measured across a field of view and divided by the number of nuclei within the same field of view. Values were normalized to the D21 average.

All transition zone antibodies were co-stained with GT335. The transition zone was identified by the gap in GT335 signal between the mother centriole and the base of the cilium. Transition zone protein intensity was determined by centering a 1 μm x 1 μm box over the transition zone and measuring the integrated density. Background was determined by a 1 μm x 1 μm box randomly placed near the centrosome. Background subtracted values were normalized to the D21 average. For TMEM67, the ROI was a 0.5 μm x 0.5 μm box to avoid centrosomal TMEM67.

### Statistical Analysis

All data sets were tested for Normality using D’Agostino and Pearson Omnibus Normality Test in GraphPad Prism 9. Normal data sets were then tested for significance with a two-tailed unpaired t-test and non-normal data sets were tested for significance using the Mann-Whitney test. All graphs show mean ± SD. All experiments utilized at least three biological replicates, unless otherwise noted.

### Electron Tomography

RPE1 cells were grown on sapphire discs and prepared for electron microscopy using high pressure freezing and freeze substitution as described in McDonald et al.^66^ Briefly, 3mm sapphire discs (Technotrade International) were coated with gold and a large F was then scratched into the surface to help orient the cell side. The disks were coated with collagen, sterilized under UV light and cells plated for culturing. Monolayers grown on sapphire discs were then frozen using a Wohlwend Compact02 high pressure freezer (Technotrade International). The frozen cells were then freeze substituted in 1% OsO4 and 0.1% uranyl acetate in acetone at −80°C for 3 days then gradually warmed to room temperature. The discs were then flat embedded in a thin layer of Epon resin and polymerized at 60°C. Regions containing cells were identified in the light microscope, and a small square of resin containing the cells was excised and remounted onto a blank resin block. The cells were then sectioned en face and serial, thick sections (250-300nm) were collected onto formvar-coated slot grids. Grids were post stained with 2% uranyl acetate and Reynold’s lead citrate and 15 nm colloidal gold (BBI International) was affixed to the section surface to serve as alignment markers.

Tomography was performed using a Tecnai F30 microscope operating at 300 kV (Thermo Fisher Scientific, Waltham, MA). Dual axis tilt series were collected over a +/- 60° range using the SerialEM image acquisition software,^67^ and a Gatan OneView camera (Gatan, Inc., Pleasanton, CA). For most data sets, tilt series were collected from 2-4 serial sections to reconstruct a larger volume of cell data. Tomograms were computed, serial tomograms joined and cellular features were modeled using the IMOD 4.9 software package (https://bio3d.colorado.edu/imod/).^68,69^

Organelles at the centrosomes of the cells (centrioles, vesicles, positions of MT and their ends) were manually traced in these reconstructions using the 3dmod program in the IMOD software package.^68^ Models were projected in 3D to show the arrangement of the centrioles, vesicles, microtubules and the position their ends within the volume. In total, 4 centrosomes from D21, 3 centrosomes from T21, and 4 centrosomes from Q21 were reconstructed and modeled. The modeled centrosomes came from different stages, including at or before ciliary vesicle formation, and one D21 and T21 set with a primary cilium.

## Supplemental Figure Legends

*Figure S1*. (A-C) Quantitation of whole cell PCNT intensities for D21 (A), T21 (B), and Q21 (C) cells throughout the time course normalized to average at 0 h. Graph shows mean ± SD. (D) Quantitation of centrosomal PCNT intensities in a 0-1.2 μm radial circle around the centrosome throughout the time course normalized to D21 average at 0 h. Graph shows mean ± SD. (E) Quantitation of pericentrosomal PCNT intensities in a 1.2-5 μm radial circle around the centrosome throughout the time course normalized to D21 average at 0 h. Graph shows mean ± SD. (F) Quantitation of centrosomal microtubule intensities in a 0-1.2 μm radial circle around the centrosome throughout the time course normalized to D21 average at 0 h. Graph shows mean ± SD. (G) Quantitation of pericentrosomal microtubule intensities in a 1.2-5 μm radial circle around the centrosome throughout the time course normalized to D21 average at 0 h. Graph shows mean ± SD. N’s and statistical tests are listed in Table S1.

*Figure S2*. (A, B) Representative SIM images of RPE1 D21, T21, and Q21 cells grown on coverslips and serum depleted for 24 h. Cells were stained with GT335 and the distal appendage marker CEP83 (A) or centrin, Acetylated tubulin (Actub), and the subdistal appendage marker Ninein (NIN) (B). Percentages represent cells with indicated marker for 3 N’s.

*Figure S3*. (A) Quantitation of whole cell MYOVA intensity for D21, T21, and Q21 cells. (B-D) Distribution of centrosomal MYOVA intensities moving away from the centrosome for 2 (B), 4 (C), and 24 (D) h timepoints. Values were normalized to the D21 average. Graph shows mean ± SD. (E) Representative SIM images of RPE1 D21, T21, and Q21 cells grown on coverslips and serum depleted for 24 h. Cells were either treated with control or PCNT siRNA during the 24 h serum depletion. Cells were stained with GT335 and MYOVA. (F) Quantitation of PCNT intensities in a 5 μm radial circle around the centrosome in control and PCNT siRNA treated cells normalized to the D21 average. Graph shows mean ± SD. (G, H) Correlation analysis between PCNT and MYOVA levels for D21, T21, and Q21 cells 24 h after serum depletion. (I) Representative confocal images of RPE1 D21, T21, and Q21 cells stably expressing GFP-EHD1 grown on coverslips and serum depleted for 24 h. Cells were stained for PCNT and Actub. N’s and statistical tests are listed in Table S1.

*Figure S4*. (A) Quantitation of CP110 centriole capping for D21, T21, and Q21 cells 24 h after serum depletion. (B) Representative SIM images of RPE1 D21, T21, and Q21 cells grown on coverslips and serum depleted for 24 h. Cells were stained with GT335 and the centriole capping protein CEP97. Percentages represent cells with indicated phenotype across 3 N’s. (C) Quantitation of CEP97 centriole capping for D21, T21, and Q21 cells. (D) Representative SIM images of RPE1 D21, T21, and Q21 cells grown on coverslips and serum depleted for 2, 4, 8, 24, and 48 h. Cells were stained with γ-tubulin, Actub, and RAB8. Percentages represent cells with indicated phenotype across 3 N’s. (E) Representative confocal images of RPE1 D21, T21, and Q21 cells grown on coverslips and serum depleted for 24 h. Cells were stained with Hoechst, PCNT, Golgin97, and RAB8. (F) Quantitation of RAB8 Golgi intensity normalized to the D21 average. (G) Cartoon model depicting intracellular trafficking pathways. (H) Representative confocal images of RPE1 D21, T21, and Q21 cells grown on coverslips and serum depleted for 24 h. Cells were stained with Hoechst, γ-tubulin, and the cis-Golgi marker GM130. (I-J) Quantitation of GM130 intensity (I) and GM130-labeled Golgi area (J) normalized to the D21 average. (K) Representative confocal images of RPE1 D21, T21, and Q21 cells grown on coverslips and serum depleted for 24 h. Cells were stained with Hoechst, PCNT, and the trans-Golgi marker Golgin97. (L-M) Quantitation of Golgin97 intensity (L) and Golgin97-labeled Golgi area (M) normalized to the D21 average. (N) Representative confocal images of RPE1 D21, T21, and Q21 cells grown on coverslips and serum depleted for 24 h. Cells were stained with Hoechst, γ-tubulin, and the early endosome marker EEA1. (O) Quantitation of EEA1 intensity normalized to the D21 average. (P) Representative confocal images of RPE1 D21, T21, and Q21 cells grown on coverslips and serum depleted for 24 h. Cells were stained with Hoechst, PCNT, and the lysosome marker CD63. Q) Quantitation of CD63 intensity normalized to the D21 average. Graphs shows mean ± SD. N’s and statistical tests are listed in Table S1.

*Figure S5*. (A) Representative confocal images of RPE1 D21 and T21 cells stably expressing pH-Smoothened (pH-SMO) grown on coverslips and serum depleted for 24 h. The IN/OUT assay was performed, and cells were stained with an anti-GFP antibody pre-permeabilization and Actub after. Anti-GFP labeled cilia are extracellular. (B) Quantitation of extracellular cilia in D21 and T21 cells. (C) Quantitation of centrosomal CEP290 intensity in 0-1.2 μm region around centrosome.

*Figure S6*. For all following experiments, WT and Dp primary MEFs were isolated from E12.5 dams, grown on coverslips, and serum depleted for 24 h. (A, B) Representative confocal images of WT and Dp16 (A) or Dp17 (B) MEFs. Cells were stained with Hoechst, the ciliary marker ARL13B, and γ-tubulin. (C, D) Representative confocal images of WT and Dp16 (C) or Dp17 (D) MEFs. Cells were stained with PCNT and ARL13B. (E) Representative confocal images of WT and the Dp10-2 MEF line that does not have decreased cilia. Cells were stained with Hoechst, the ciliary marker ARL13B, and Actub. (F) Quantitation of the number of cells with a primary cilium in WT and Dp10-2 MEFs. (G) Quantitation of PCNT intensities in a 5 μm radial circle around the centrosome in WT and Dp10-2 MEFs. Values were normalized to the WT average. (H) Quantitation of ciliary SMO levels in SAG-treated cells for WT and Dp10-2 MEFs. Values were normalized to the WT average. Graphs show mean ± SD. N’s and statistical tests are listed in Table S1.

*Figure S7*. (A) Representative tiled confocal images of WT and Dp10 P4 animals corresponding to the same cerebellar folia in each animal. Brain sections were stained with Hoechst and the cell proliferation marker Ki67. (B) Quantitation of the number of Ki67-positive cells in the external granular layer of WT and Dp10 animals. (C) Representative tiled confocal images of WT and Dp10 P4 animals corresponding to the same cerebellar folia in each animal. Brain sections were stained with Hoechst and the Purkinje cell marker calbindin (CALB1) and the microtubule binding protein doublecortin (DCX). The far-right panel shows insets of DCX staining. (D-F) Representative tiled confocal images of multiciliated ependymal cells lining the ventricle next to the cerebellum from P4 wild-type (WT) and Dp10 (E), Dp16 (F), and Dp17 (G) animals. Brain sections were stained with Hoechst, ARL13B, and γ-tubulin.

*Video S1-3*. Movies of D21 (S1), T21 (S2), and Q21 (S3) cells from Figure 2 showing the EM tomogram volume, then models projecting from the images, then the model turning without images. Mother centriole (yellow), daughter centriole (magenta), microtubule minus ends (light blue spheres), microtubules (green), and vesicles (red). Note the proximity of vesicles to microtubules.

## References

1 Haydar, T. F. & Reeves, R. H. Trisomy 21 and early brain development. Trends Neurosci 35, 81–91, doi:10.1016/j.tins.2011.11.001 (2012).

2 Bergström, S. et al. Trends in Congenital Heart Defects in Infants With Down Syndrome. Pediatrics 138, doi:10.1542/peds.2016-0123 (2016).

3 Richtsmeier, J. T., Baxter, L. L. & Reeves, R. H. Parallels of craniofacial maldevelopment in Down syndrome and Ts65Dn mice. Developmental dynamics: an official publication of the American Association of Anatomists 217, 137–145, doi:10.1002/(sici)1097-0177(200002)217:2<137::Aid-dvdy1>3.0.Co;2-n (2000).

4 Goetz, S. C. & Anderson, K. V. The primary cilium: a signalling centre during vertebrate development. Nat Rev Genet 11, 331–344, doi:10.1038/nrg2774 (2010).

5 Schmidt, K. N. et al. Cep164 mediates vesicular docking to the mother centriole during early steps of ciliogenesis. J Cell Biol 199, 1083–1101, doi:10.1083/jcb.201202126 (2012).

6 Tanos, B. E. et al. Centriole distal appendages promote membrane docking, leading to cilia initiation. Genes Dev 27, 163–168, doi:10.1101/gad.207043.112 (2013).

7 Ishikawa, H., Kubo, A., Tsukita, S. & Tsukita, S. Odf2-deficient mother centrioles lack distal/subdistal appendages and the ability to generate primary cilia. Nat Cell Biol 7, 517–524, doi:10.1038/ncb1251 (2005).

8 Wu, C. T., Chen, H. Y. & Tang, T. K. Myosin-Va is required for preciliary vesicle transportation to the mother centriole during ciliogenesis. Nat Cell Biol 20, 175–185, doi:10.1038/s41556-017-0018-7 (2018).

9 Lu, Q. et al. Early steps in primary cilium assembly require EHD1/EHD3-dependent ciliary vesicle formation. Nat Cell Biol 17, 228–240, doi:10.1038/ncb3109 (2015).

10 Garcia-Gonzalo, F. R. & Reiter, J. F. Scoring a backstage pass: mechanisms of ciliogenesis and ciliary access. J Cell Biol 197, 697–709, doi:10.1083/jcb.201111146 (2012).

11 Sung, C. H. & Leroux, M. R. The roles of evolutionarily conserved functional modules in cilia-related trafficking. Nat Cell Biol 15, 1387–1397, doi:10.1038/ncb2888 (2013).

12 Nachury, M. V., Seeley, E. S. & Jin, H. Trafficking to the ciliary membrane: how to get across the periciliary diffusion barrier? Annu Rev Cell Dev Biol 26, 59–87, doi:10.1146/annurev.cellbio.042308.113337 (2010).

13 Kim, J., Krishnaswami, S. R. & Gleeson, J. G. CEP290 interacts with the centriolar satellite component PCM-1 and is required for Rab8 localization to the primary cilium. Hum Mol Genet 17, 3796–3805, doi:10.1093/hmg/ddn277 (2008).

14 Blacque, O. E., Scheidel, N. & Kuhns, S. Rab GTPases in cilium formation and function. Small GTPases 9, 76–94, doi:10.1080/21541248.2017.1353847 (2018).

15 Hori, A. & Toda, T. Regulation of centriolar satellite integrity and its physiology. Cell Mol Life Sci 74, 213–229, doi:10.1007/s00018-016-2315-x (2017).

16 Doxsey, S. J., Stein, P., Evans, L., Calarco, P. D. & Kirschner, M. Pericentrin, a highly conserved centrosome protein involved in microtubule organization. Cell 76, 639–650, doi:10.1016/0092-8674(94)90504-5 (1994).

17 Gavilan, M. P. et al. The dual role of the centrosome in organizing the microtubule network in interphase. EMBO Rep 19, doi:10.15252/embr.201845942 (2018).

18 Martinez-Campos, M., Basto, R., Baker, J., Kernan, M. & Raff, J. W. The Drosophila pericentrin-like protein is essential for cilia/flagella function, but appears to be dispensable for mitosis. J Cell Biol 165, 673–683, doi:10.1083/jcb.200402130 (2004).

19 Galati, D. F., Sullivan, K. D., Pham, A. T., Espinosa, J. M. & Pearson, C. G. Trisomy 21 Represses Cilia Formation and Function. Dev Cell 46, 641–650.e646, doi:10.1016/j.devcel.2018.07.008 (2018).

20 McCurdy, B. L. et al. Trisomy 21 increases microtubules and disrupts centriolar satellite localization. bioRxiv (2021).

21 Joo, K. et al. CCDC41 is required for ciliary vesicle docking to the mother centriole. Proc Natl Acad Sci U S A 110, 5987–5992, doi:10.1073/pnas.1220927110 (2013).

22 Shakya, S. & Westlake, C. J. Recent advances in understanding assembly of the primary cilium membrane. Fac Rev 10, 16, doi:10.12703/r/10-16 (2021).

23 Yang, T. T. et al. Super-resolution architecture of mammalian centriole distal appendages reveals distinct blade and matrix functional components. Nat Commun 9, 2023, doi:10.1038/s41467-018-04469-1 (2018).

24 Lange, B. M. & Gull, K. A molecular marker for centriole maturation in the mammalian cell cycle. J Cell Biol 130, 919–927, doi:10.1083/jcb.130.4.919 (1995).

25 Nakagawa, Y., Yamane, Y., Okanoue, T., Tsukita, S. & Tsukita, S. Outer dense fiber 2 is a widespread centrosome scaffold component preferentially associated with mother centrioles: its identification from isolated centrosomes. Mol Biol Cell 12, 1687–1697, doi:10.1091/mbc.12.6.1687 (2001).

26 Mogensen, M. M., Malik, A., Piel, M., Bouckson-Castaing, V. & Bornens, M. Microtubule minus-end anchorage at centrosomal and non-centrosomal sites: the role of ninein. J Cell Sci 113 (Pt 17), 3013–3023 (2000).

27 Spektor, A., Tsang, W. Y., Khoo, D. & Dynlacht, B. D. Cep97 and CP110 suppress a cilia assembly program. Cell 130, 678–690, doi:10.1016/j.cell.2007.06.027 (2007).

28 Liu, M. et al. NudCL2 is an autophagy receptor that mediates selective autophagic degradation of CP110 at mother centrioles to promote ciliogenesis. Cell Res, doi:10.1038/s41422-021-00560-3 (2021).

29 Nachury, M. V. et al. A Core Complex of BBS Proteins Cooperates with the GTPase Rab8 to Promote Ciliary Membrane Biogenesis. Cell 129, 1201–1213, doi:10.1016/j.cell.2007.03.053 (2007).

30 Moritz, O. L. et al. Mutant rab8 Impairs docking and fusion of rhodopsin-bearing postGolgi membranes and causes cell death of transgenic Xenopus rods. Mol Biol Cell 12, 2341–2351, doi:10.1091/mbc.12.8.2341 (2001).

31 Sorokin, S. CENTRIOLES AND THE FORMATION OF RUDIMENTARY CILIA BY FIBROBLASTS AND SMOOTH MUSCLE CELLS. J Cell Biol 15, 363, doi:10.1083/jcb.15.2.363 (1962).

32 Kukic, I., Rivera-Molina, F. & Toomre, D. The IN/OUT assay: a new tool to study ciliogenesis. Cilia 5, 23, doi:10.1186/s13630-016-0044-2 (2016).

33 Tobin, J. L. & Beales, P. L. The nonmotile ciliopathies. Genet Med 11, 386–402, doi:10.1097/GIM.0b013e3181a02882 (2009).

34 Sang, L. et al. Mapping the NPHP-JBTS-MKS protein network reveals ciliopathy disease genes and pathways. Cell 145, 513–528, doi:10.1016/j.cell.2011.04.019 (2011).

35 Craige, B. et al. CEP290 tethers flagellar transition zone microtubules to the membrane and regulates flagellar protein content. J Cell Biol 190, 927–940, doi:10.1083/jcb.201006105 (2010).

36 Yang, T. T. et al. Superresolution Pattern Recognition Reveals the Architectural Map of the Ciliary Transition Zone. Sci Rep 5, 14096, doi:10.1038/srep14096 (2015).

37 Williams, C. L. et al. MKS and NPHP modules cooperate to establish basal body/transition zone membrane associations and ciliary gate function during ciliogenesis. J Cell Biol 192, 1023–1041, doi:10.1083/jcb.201012116 (2011).

38 Garcia-Gonzalo, F. R. et al. A transition zone complex regulates mammalian ciliogenesis and ciliary membrane composition. Nat Genet 43, 776–784, doi:10.1038/ng.891 (2011).

39 Roper, R. J. et al. Defective cerebellar response to mitogenic Hedgehog signaling in Down [corrected] syndrome mice. Proc Natl Acad Sci U S A 103, 1452–1456, doi:10.1073/pnas.0510750103 (2006).

40 Gupta, M., Dhanasekaran, A. R. & Gardiner, K. J. Mouse models of Down syndrome: gene content and consequences. Mamm Genome 27, 538–555, doi:10.1007/s00335-016-9661-8 (2016).

41 Yu, T. et al. Effects of individual segmental trisomies of human chromosome 21 syntenic regions on hippocampal long-term potentiation and cognitive behaviors in mice. Brain Research 1366, 162–171, doi:10.1016/j.brainres.2010.09.107 (2010).

42 van Dam, T. J., Wheway, G., Slaats, G. G., Huynen, M. A. & Giles, R. H. The SYSCILIA gold standard (SCGSv1) of known ciliary components and its applications within a systems biology consortium. Cilia 2, 7, doi:10.1186/2046-2530-2-7 (2013).

43 Kong, J. H., Siebold, C. & Rohatgi, R. Biochemical mechanisms of vertebrate hedgehog signaling. Development 146, doi:10.1242/dev.166892 (2019).

44 Ho, E. K. & Stearns, T. Hedgehog signaling and the primary cilium: implications for spatial and temporal constraints on signaling. Development 148, doi:10.1242/dev.195552 (2021).

45 Smeyne, R. J. et al. Local control of granule cell generation by cerebellar Purkinje cells. Mol Cell Neurosci 6, 230–251, doi:10.1006/mcne.1995.1019 (1995).

46 Wechsler-Reya, R. J. & Scott, M. P. Control of neuronal precursor proliferation in the cerebellum by Sonic Hedgehog. Neuron 22, 103–114, doi:10.1016/s0896-6273(00)80682-0 (1999).

47 Spassky, N. et al. Primary cilia are required for cerebellar development and Shh-dependent expansion of progenitor pool. Dev Biol 317, 246–259, doi:10.1016/j.ydbio.2008.02.026 (2008).

48 Haldipur, P. et al. Preterm delivery disrupts the developmental program of the cerebellum. PLOS ONE 6, e23449, doi:10.1371/journal.pone.0023449 (2011).

49 Nguyen, V. et al. Sonic Hedgehog Agonist Protects Against Complex Neonatal Cerebellar Injury. Cerebellum 17, 213–227, doi:10.1007/s12311-017-0895-0 (2018).

50 Delaval, B. & Doxsey, S. J. Pericentrin in cellular function and disease. J Cell Biol 188, 181–190, doi:10.1083/jcb.200908114 (2010).

51 Endoh-Yamagami, S. et al. A mutation in the pericentrin gene causes abnormal interneuron migration to the olfactory bulb in mice. Dev Biol 340, 41–53, doi:10.1016/j.ydbio.2010.01.017 (2010).

52 Gleeson, J. G., Lin, P. T., Flanagan, L. A. & Walsh, C. A. Doublecortin is a microtubule-associated protein and is expressed widely by migrating neurons. Neuron 23, 257–271, doi:10.1016/s0896-6273(00)80778-3 (1999).

53 Francis, F. et al. Doublecortin is a developmentally regulated, microtubule-associated protein expressed in migrating and differentiating neurons. Neuron 23, 247–256, doi:10.1016/s0896-6273(00)80777-1 (1999).

54 Nishita, M., Satake, T., Minami, Y. & Suzuki, A. Regulatory mechanisms and cellular functions of non-centrosomal microtubules. J Biochem 162, 1–10, doi:10.1093/jb/mvx018 (2017).

55 Hibbard, J. V. K., Vazquez, N., Satija, R. & Wallingford, J. B. Protein turnover dynamics suggest a diffusion-to-capture mechanism for peri-basal body recruitment and retention of intraflagellar transport proteins. Mol Biol Cell, mbcE20110717, doi:10.1091/mbc.E20-11-0717 (2021).

56 Duchon, A. et al. Identification of the translocation breakpoints in the Ts65Dn and Ts1Cje mouse lines: relevance for modeling Down syndrome. Mamm Genome 22, 674–684, doi:10.1007/s00335-011-9356-0 (2011).

57 Stemm-Wolf, A. J., O’Toole, E. T., Sheridan, R. M., Morgan, J. T. & Pearson, C. G. The SON RNA splicing factor is required for intracellular trafficking structures that promote centriole assembly and ciliogenesis. Mol Biol Cell, mbcE21060305, doi:10.1091/mbc.E21-06-0305 (2021).

58 Ahn, E. Y. et al. SON controls cell-cycle progression by coordinated regulation of RNA splicing. Mol Cell 42, 185–198, doi:10.1016/j.molcel.2011.03.014 (2011).

59 Yu, T. et al. A mouse model of Down syndrome trisomic for all human chromosome 21 syntenic regions. Hum Mol Genet 19, 2780–2791, doi:10.1093/hmg/ddq179 (2010).

60 Del Viso, F. et al. Congenital Heart Disease Genetics Uncovers Context-Dependent Organization and Function of Nucleoporins at Cilia. Dev Cell 38, 478–492, doi:10.1016/j.devcel.2016.08.002 (2016).

61 Lane, A. A. et al. Triplication of a 21q22 region contributes to B cell transformation through HMGN1 overexpression and loss of histone H3 Lys27 trimethylation. Nat Genet 46, 618–623, doi:10.1038/ng.2949 (2014).

62 Mariani, L. E. et al. Arl13b regulates Shh signaling from both inside and outside the cilium. Mol Biol Cell 27, 3780–3790, doi:10.1091/mbc.E16-03-0189 (2016).

63 Waterman-Storer, C. M. & Salmon, E. D. Actomyosin-based retrograde flow of microtubules in the lamella of migrating epithelial cells influences microtubule dynamic instability and turnover and is associated with microtubule breakage and treadmilling. J Cell Biol 139, 417–434, doi:10.1083/jcb.139.2.417 (1997).

64 Schindelin, J. et al. Fiji: an open-source platform for biological-image analysis. Nat Methods 9, 676–682, doi:10.1038/nmeth.2019 (2012).

65 Liu, Y. et al. Myelin-specific multiple sclerosis antibodies cause complement-dependent oligodendrocyte loss and demyelination. Acta Neuropathol Commun 5, 25, doi:10.1186/s40478-017-0428-6 (2017).

66 McDonald, K. et al. “Tips and tricks” for high-pressure freezing of model systems. Methods in cell biology 96, 671–693, doi:10.1016/s0091-679x(10)96028-7 (2010).

67 Mastronarde, D. N. Automated electron microscope tomography using robust prediction of specimen movements. J Struct Biol 152, 36–51, doi:10.1016/j.jsb.2005.07.007 (2005).

68 Kremer, J. R., Mastronarde, D. N. & McIntosh, J. R. Computer visualization of three-dimensional image data using IMOD. J Struct Biol 116, 71–76, doi:10.1006/jsbi.1996.0013 (1996).

69 Mastronarde, D. N. Dual-axis tomography: an approach with alignment methods that preserve resolution. J Struct Biol 120, 343–352, doi:10.1006/jsbi.1997.3919 (1997).

