## Supplementary figures and images for "Trisomy 21 induces pericentrosomal crowding disrupting early stages of primary ciliogenesis and mouse cerebellar development"

### Supplemental Figure 1

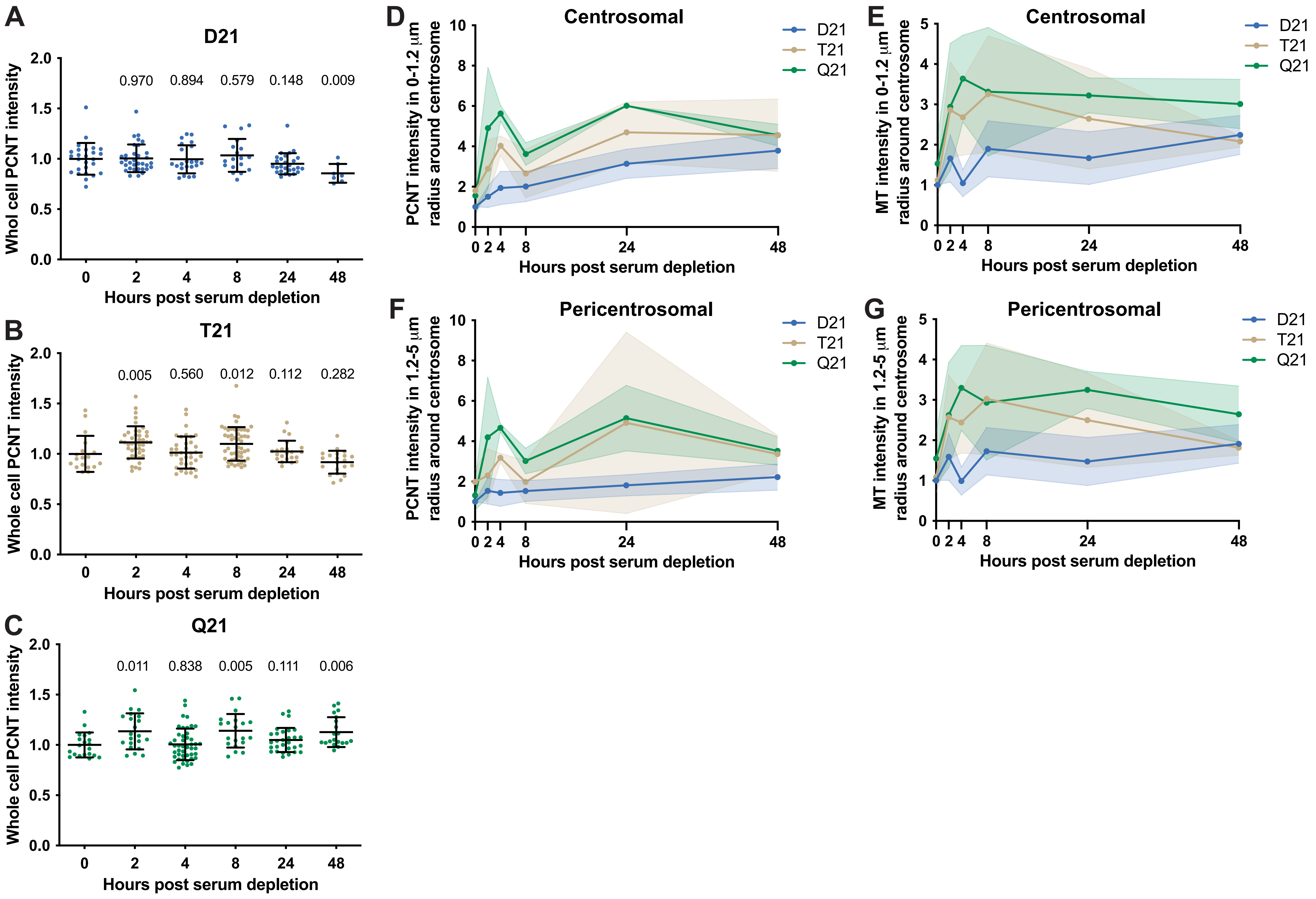

### Supplemental Figure 2

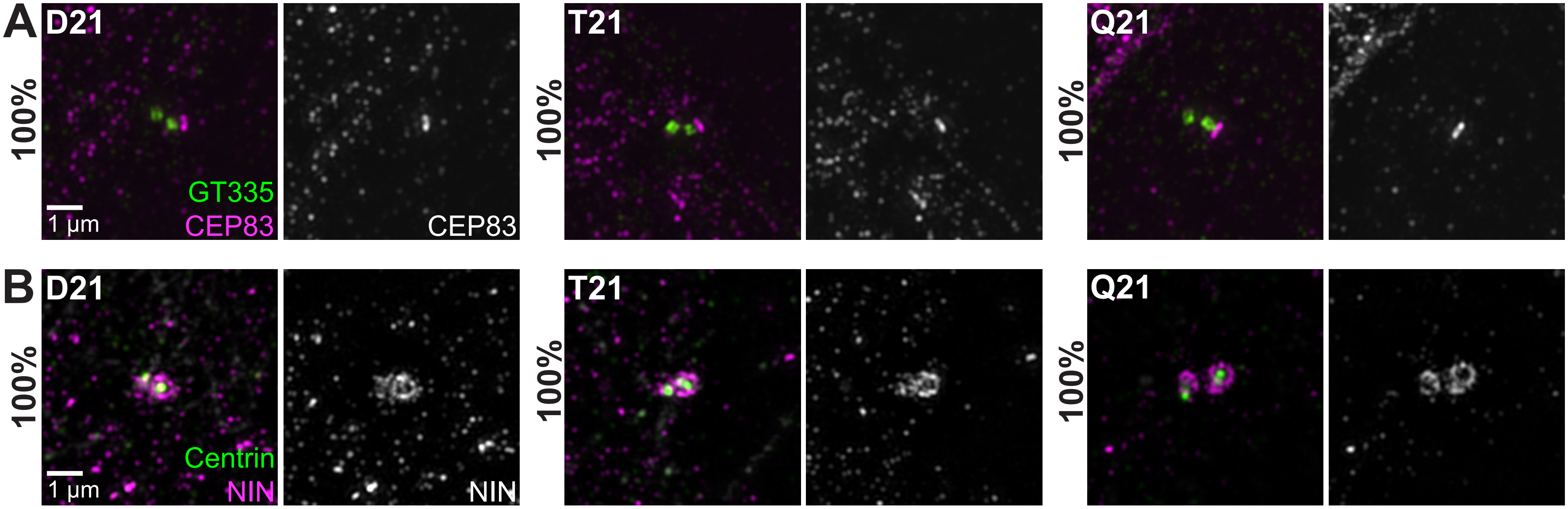

### Supplemental Figure 3

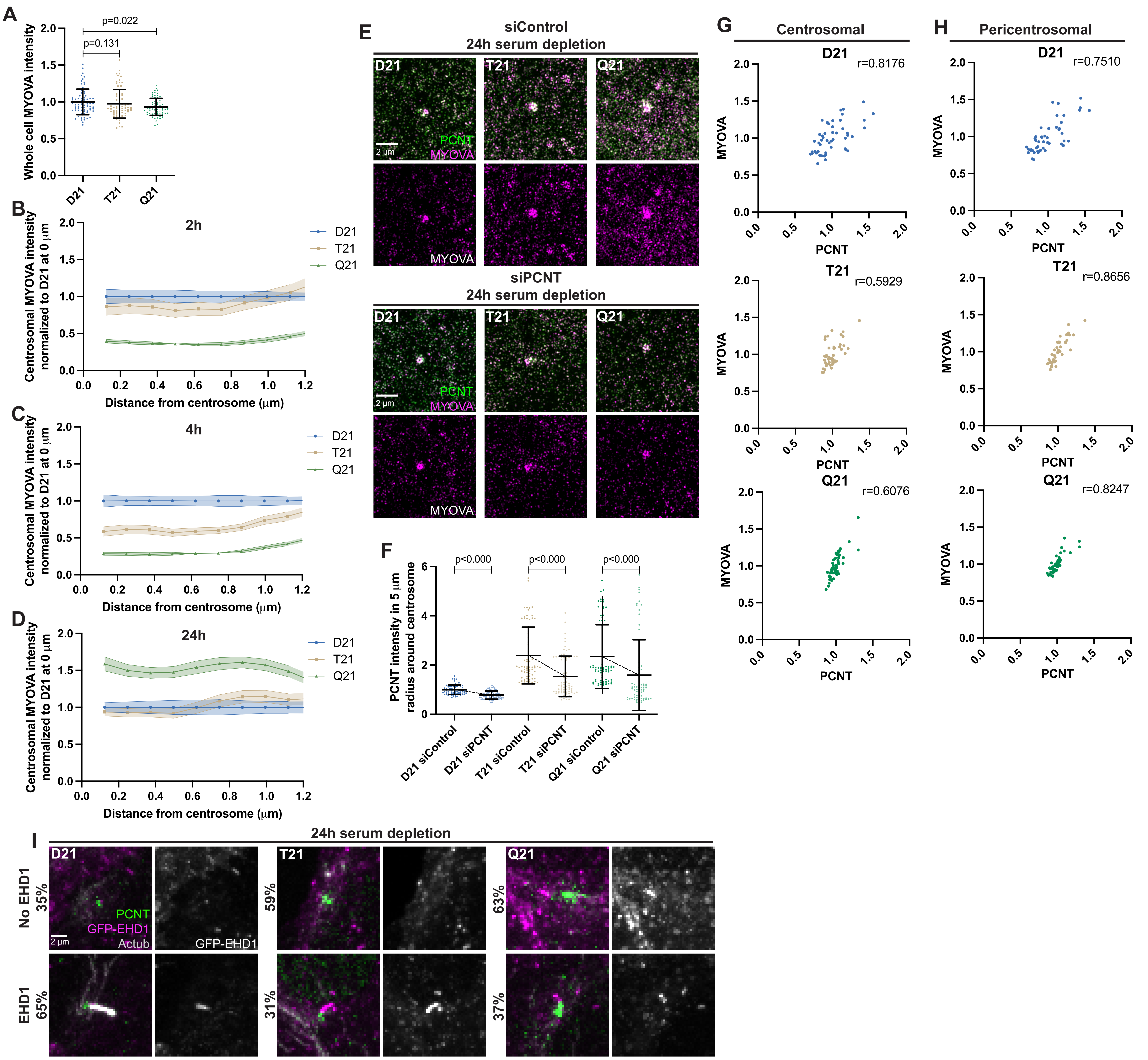

### Supplemental Figure 4

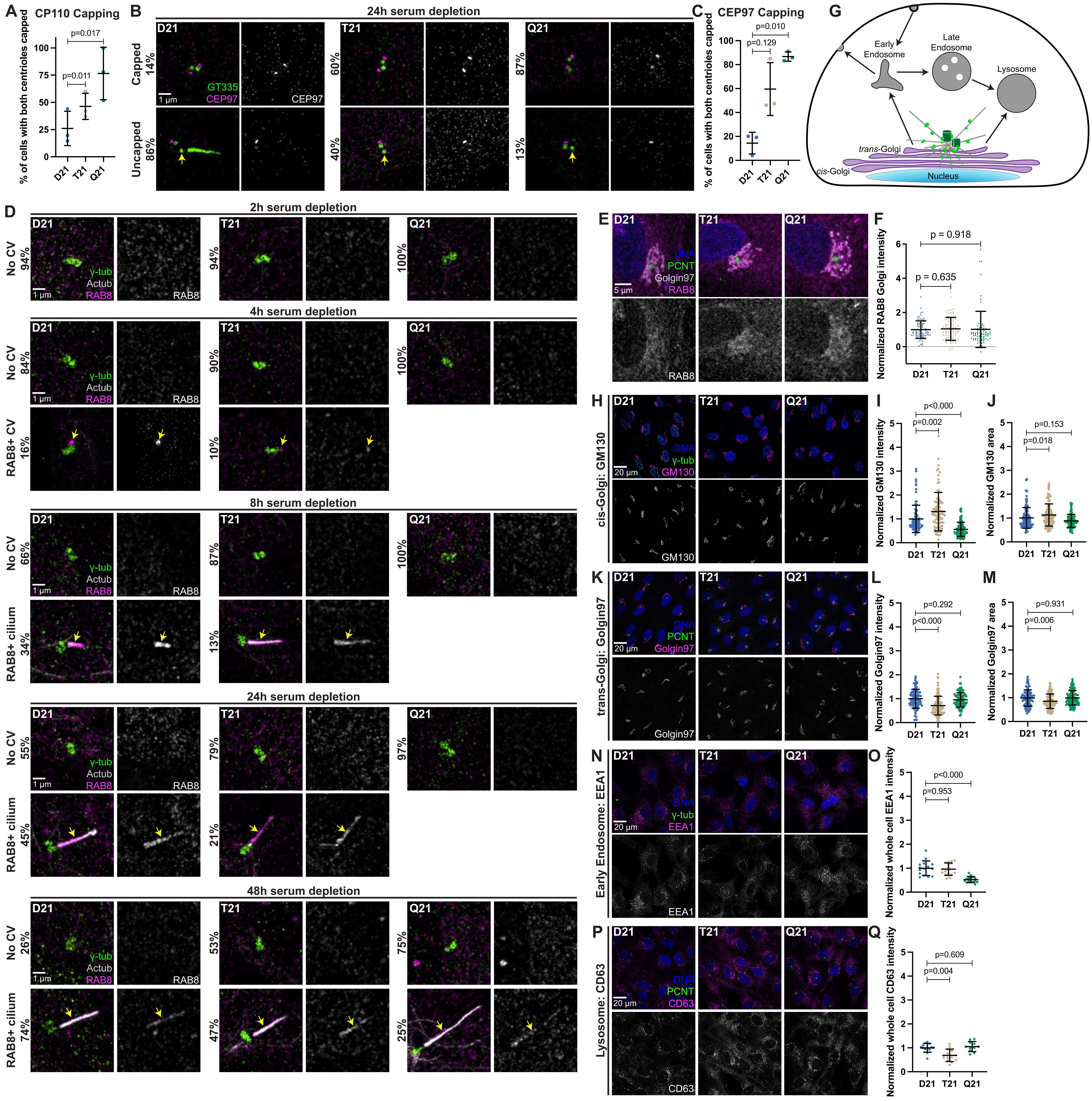

### Supplemental Figure 5

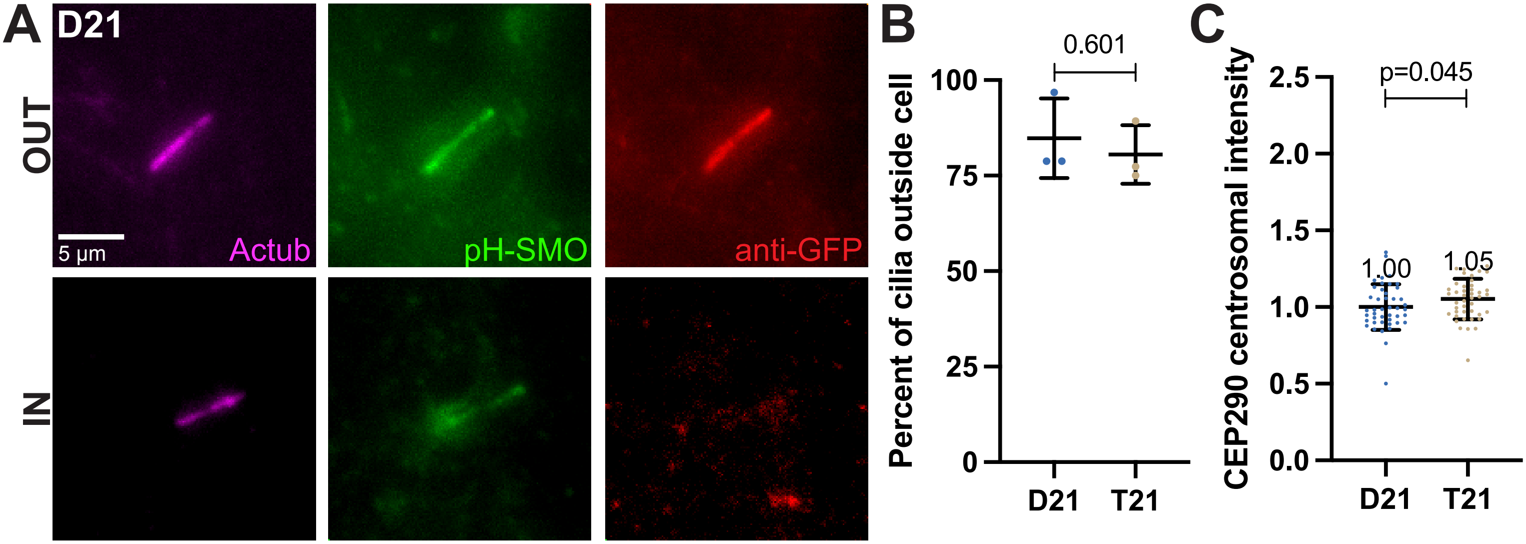

### Supplemental Figure 6

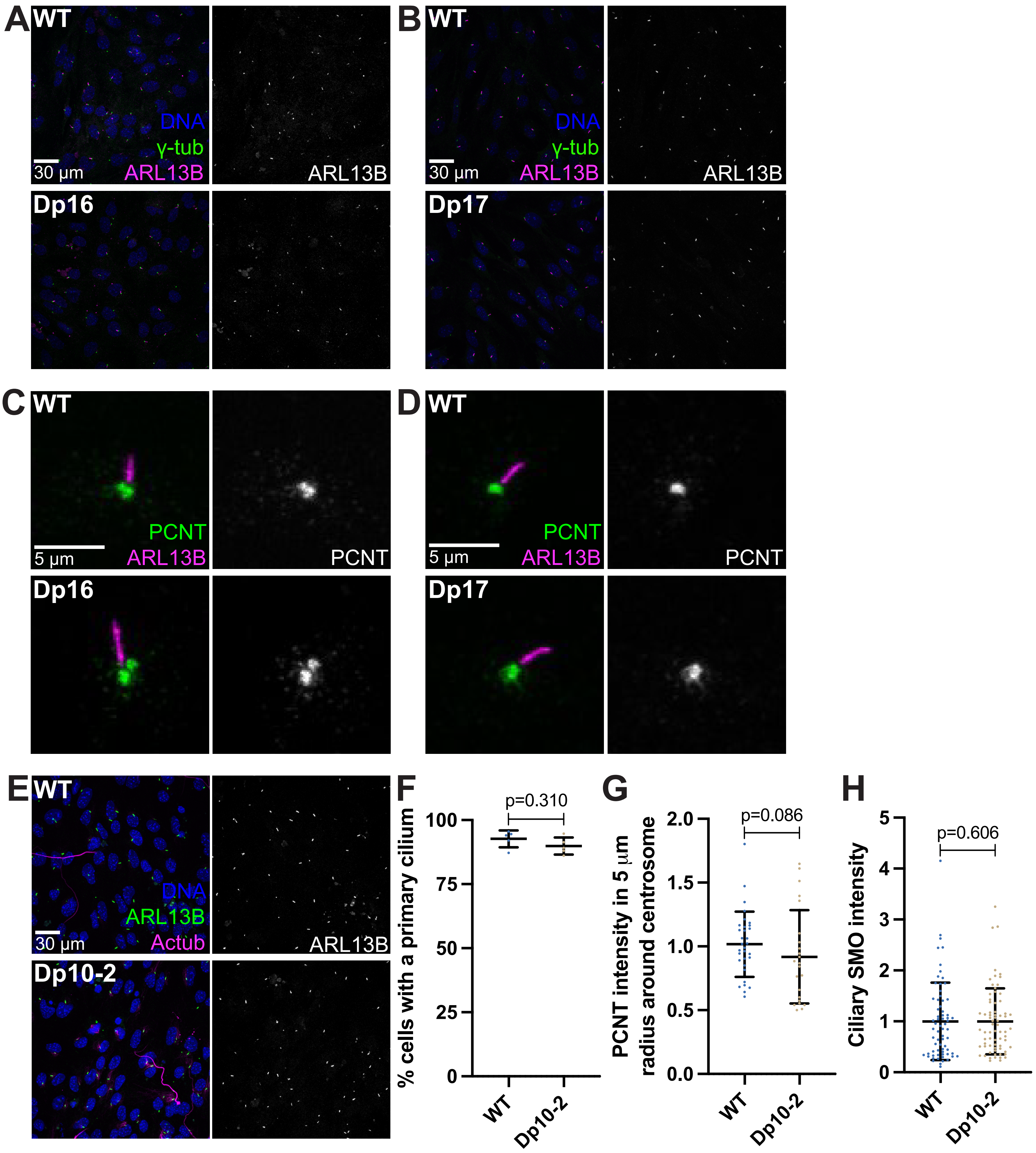

### Supplemental Figure 7

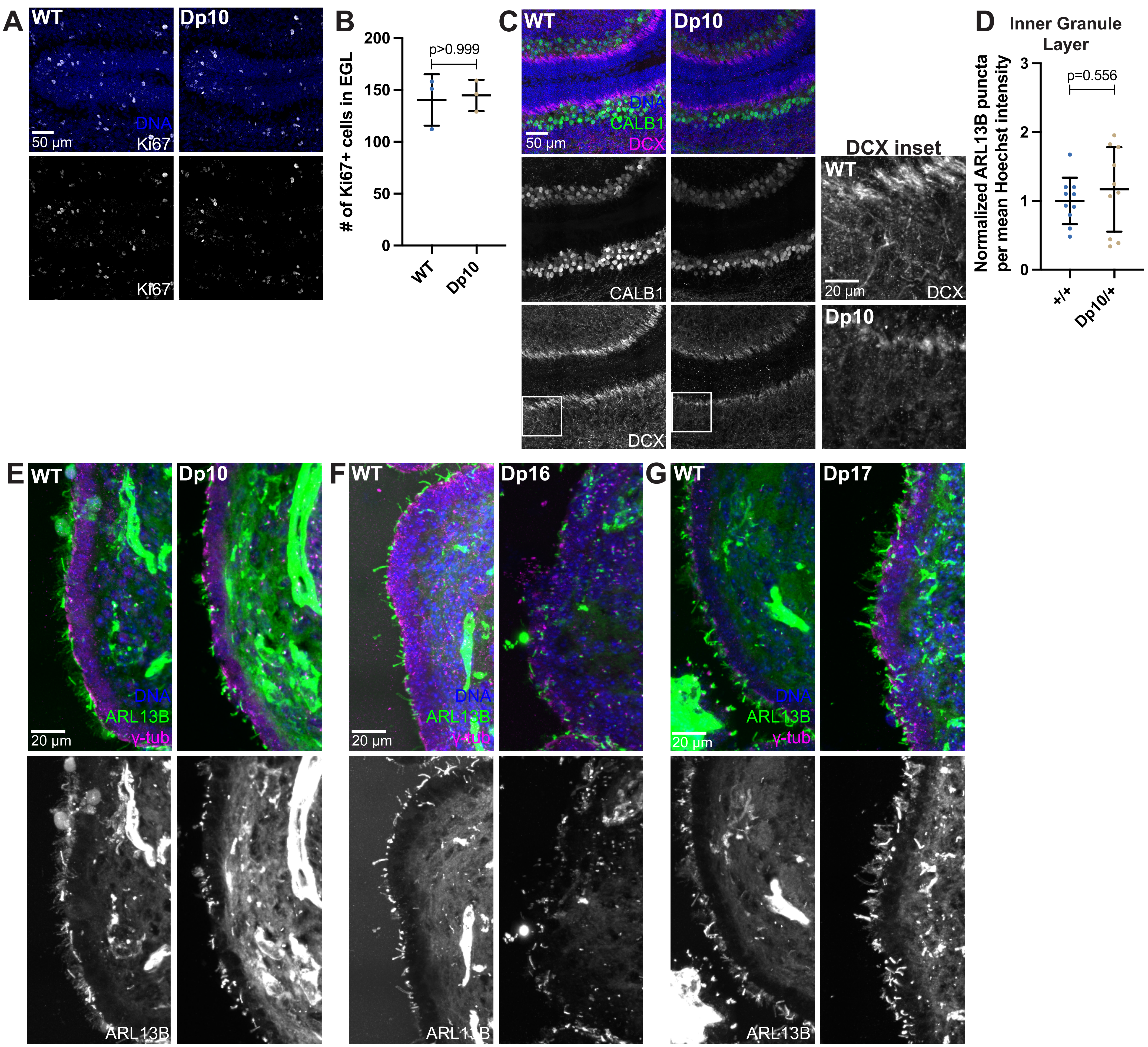
